# Dynamic embedding of salience coding in hippocampal spatial maps

**DOI:** 10.1101/266767

**Authors:** Masaaki Sato, Kotaro Mizuta, Tanvir Islam, Masako Kawano, Takashi Takekawa, Daniel Gomez-Dominguez, Karam Kim, Hiroshi Yamakawa, Masamichi Ohkura, Tomoki Fukai, Junichi Nakai, Yasunori Hayashi

## Abstract

Hippocampal CA1 neurons participate in dynamic ensemble codes for space and memory. Prominent features of the environment are represented by an increased density of place cells, but cellular principles governing the formation and plasticity of such disproportionate maps are unknown. We thus imaged experience-dependent long-term changes in spatial representations at the cellular level in the CA1 deep sublayer in mice learning to navigate in a virtual-reality environment. The maps were highly dynamic but gradually stabilized as over-representations for motivational (reward) and environmental (landmark) salience emerged in different time courses by selective consolidation of relevant spatial representations. Relocation of the reward extensively reorganized pre-formed maps by a mechanism involving rapid recruitment of cells from the previous location followed by their re-stabilization, indicating that a subset of neurons encode reward-related information. The distinct properties of these CA1 cells may provide a substrate by which salient experience forms lasting and adaptable memory traces.

## Introduction

Navigation and spatial memory are essential elements of animal behavior that allow animals to forage, return home and avoid dangers. The hippocampus plays a crucial role in these cognitive processes, as hippocampal neurons fire when an animal is located in a particular part of an environment but not in others, providing an allocentric cognitive map of space (O’Keefe and Nadel, 1978). Although whether these “place cells” are indeed memory cells has been long debated, accumulating evidence supports this notion. One line of such evidence includes the knowledge that hippocampal place-specific firing exhibits dynamic changes according to context and experience on multiple time scales, ranging from a few minutes to days or weeks (Muller and Kubie, 1987; Bostock et al., 1991; Mehta et al., 1997; Lever et al., 2002; Leutgeb et al., 2005a). Moreover, studies have reported that a disproportionately large number of place cells are recorded at locations that are associated with reward, safety or walls or edges of the environment (O’Keefe and Conway, 1978; Wiener et al., 1989; Hetherington and Shapiro, 1997; Hollup et al., 2001; Dombeck et al., 2010; Dupret et al., 2010; Danielson et al., 2016), indicating that the environment surrounding an animal is not represented uniformly in the hippocampal map; their representations are strongly influenced by the motivational and environmental salience of the locations.

These findings imply that an increased number of place cells encode the presence of salience (i.e., something that draws attention) in the hippocampal map. This idea further proposes potential roles of such salience maps in not only spatial (Hollup et al., 2001; Dupret et al., 2010) or episodic-like memories (Komorowski et al., 2009; Eichenbaum and Cohen, 2014) but also goal-directed and landmark-based navigation (Gothard et al., 1996) because they can signal the subject’s distance and direction relative to the represented salience by increases and decreases of population output activity to downstream neurons (Burgess and O’Keefe, 1996). Place cells are formed rapidly within minutes after initial exposure to a new environment (Hill, 1978; Wilson and McNaughton, 1993; Frank et al., 2004). However, how the over-representation of salience is established and updated by experience has been poorly explored to date. Furthermore, mechanistic insights based on session-by-session comparisons of large-scale neuronal population data are also scarce. A few different but not mutually exclusive schemes are possible to answer this question. First, place cells are formed more preferentially for salient locations than non-salient locations from the beginning of place map formation (hereafter called the “direct formation” model) (Figure 1A). This model assumes that the probabilities of place cell formation are higher at salient locations. Second, place cells are initially formed uniformly for all locations, but cells that represent salient locations subsequently increase by recruiting the place cells that represent non-salient locations (“lateral recruitment” model) (Figure 1B). This model assumes that the probabilities of place cell formation are uniform across all locations and that the probabilities that place cells encoding non-salient locations turn into those encoding salient locations are higher than the probabilities that they continue to encode non-salient locations. Third, the place fields of all place cells potentially turn over dynamically, but spatial representations of salient locations are more stable than nonsalient locations, leading to the persistence or accumulation of relevant spatial representations over time (“selective consolidation” model) (Figure 1C).

**Figure 1.**
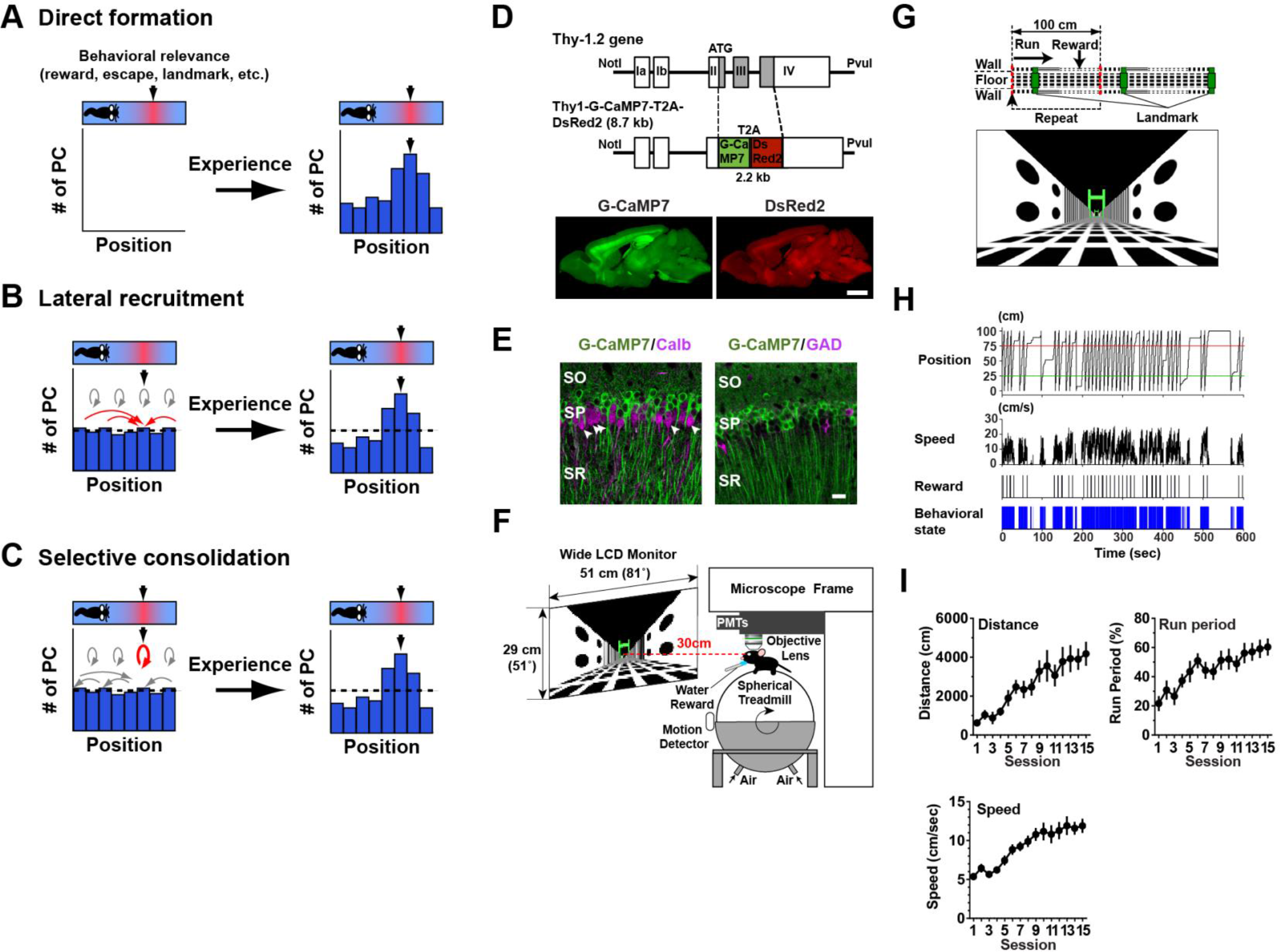
Models, transgenic mice and behavioral task. (A-C) Models that can account for the formation of hippocampal over-representation of salient locations. (A) Direct formation model. (B) Lateral recruitment model. (C) Selective consolidation model. See Introduction for details. (D) Transgene construct for Thy1-G-CaMP7 mice (top) and expression of G-CaMP7 (bottom left, green) and DsRed2 (bottom right, red) in a parasagittal section of a mouse at 6 months of age. Scale bar = 2 mm. (E) Characterization of the G-CaMP7-expressing cell population in the dorsal CA1 hippocampus of Thy1-G-CaMP7 transgenic mice. Sections of mice at 2-3 months of age were immunolabeled with anti-calbindin (Calb, left) or anti-glutamic acid decarboxylase 65/67 (GAD, right) antibodies (magenta). Arrowheads indicate examples of calbindin-positive, G-CaMP7-negative cells. SO, *stratum oriens*; SP, *stratum pyramidale*; SR, *stratum radiatum*. Scale bar = 20 μm. (F) A schematic representation of the two-photon microscope and virtual reality setup used in this study. (G) Virtual endless linear track task. The linear track segment contained a green gate as a visual landmark and a reward delivery point at two distinct locations. When the mouse’s virtual position reached the point indicated by the red dotted line in the middle, it returned to the origin, so the same track segment was presented repeatedly. The bottom panel shows a camera view of the track displayed on the LCD monitor. (H) Example behavioral data from a single 10-min session. From top to bottom, the mouse’s virtual position on the linear track, running speed, timing of reward delivery, and behavioral state during which a period of running is represented in blue. (I) Behavioral changes induced by repeated training. Total distance traveled (Distance, upper left), the fraction of time spent running (Run period, upper right) and running speed (Speed, lower left) are shown.

To elucidate the single or multiple forms of cellular dynamics that govern the formation and plasticity of the salience representation in the hippocampus, we longitudinally imaged place maps of the CA1 deep sublayer in a neuron-specific G-CaMP7 transgenic mouse line during training on a virtual linear track, in which two distinct locations were associated with either reward or a visual landmark as motivational or environmental salience, respectively. The experiments allowed us to track the neuronal activities and anatomical positions of a large population of pyramidal cells within a particular area of the hippocampus over days and thus to examine the dynamic cellular changes predicted differently by each of the above hypotheses. We found that initial establishment of over-represented salience maps is dominated by selective consolidation of the cells that encode each salient location but not by direct formation from non-place cells or lateral recruitment of the cells that encode non-salient locations. We also found that over-representations of motivational and environmental salience emerge with experience in different rapid versus delayed time courses, respectively. By contrast, robust reorganization of pre-established maps by rearrangement of salient features occurred via a coordinated interplay of these three processes. These findings reveal the distinct engagement of multiple forms of cellular dynamics in the establishment and reorganization of hippocampal salience maps and provide a mechanism by which experience of salient environmental features form lasting and adaptable traces in these maps.

## Results

### Mice and behavioral task

To reliably perform longitudinal imaging of large-scale hippocampal functional cellular maps, we generated a new transgenic mouse line that coexpresses the fluorescence calcium indicator protein G-CaMP7 and the calcium-insensitive red fluorescence marker protein DsRed2 via 2A peptide-mediated bicistronic expression under the neuron-specific Thy1 promoter. G-CaMP7 is an improved, highly sensitive G-CaMP variant that exhibits large fluorescence changes and rapid kinetics in response to a wide range of intracellular calcium concentrations (Ohkura et al., 2012; Poder et al., 2015). We selected one mouse line, here termed Thy1-G-CaMP7, that expresses G-CaMP7 and DsRed2 in widespread brain areas at a high level in the adult brain (Figure 1D). In the dorsal CA1 hippocampus, the population of calbindin-negative pyramidal cells in the deep pyramidal cell sublayer was preferentially labeled with G-CaMP7 (Mizuseki et al., 2011; Kohara et al., 2014; Lee et al., 2014; Valero et al., 2015; Danielson et al., 2016) (Figure 1E). Immunofluorescence labeling against glutamic acid decarboxylase 65/67, parvalbumin or somatostatin revealed that interneurons positive for these markers were devoid of G-CaMP7 expression (Figure 1E and S1A). In addition to strong hippocampal expression (Figure S1B), G-CaMP7 expression was found in diverse brain areas, including the cerebral cortex, olfactory bulb, brainstem and cerebellum (Figure S1C-N), making Thy1-G-CaMP7 mice an attractive alternative to our previously reported TRE-G-CaMP7 mice in various imaging studies (Sato et al., 2015).

To allow for imaging of hippocampal maps during repeated training of spatial behavior in a controlled environment, we used a head-fixed virtual reality (VR) system that consisted of an air-supported Styrofoam treadmill and a wide LCD monitor placed under a two-photon microscope (Sato et al., 2017) (Figure 1F). Thy1-G-CaMP7 mice were trained to run along a virtual linear track for water reinforcement (Figure 1G). In this task, a head-fixed mouse in a virtual environment ran unidirectionally through an open-ended linear track segment whose walls were textured with different patterns. The mouse started running from the origin of the segment, passed under a green gate as a visual landmark, received water at a reward point and returned to the origin after reaching the other end (for details, see Methods). The visual landmark and reward delivery were associated with two distinct locations in the track to examine the effects of two different kinds of salience separately. Because the transition was instantaneous and the patterns of the walls and floor appeared seamless, the mice kept running forward as if they ran along an infinitely long repetition of a corridor, similar to a real-world circular track and a head-fixed treadmill belt used elsewhere (Hollup et al., 2001; Danielson et al., 2016). In training, behavioral performance measured by distance traveled and time spent running during 10-min sessions markedly increased as training proceeded (Figure 1H-I; Distance, p < 0.0001, F_(14,154)_ = 7.30; Run period, p < 0.0001, F_(14,154)_ = 5.26; n = 12 mice from 3 groups, one-way ANOVA). In addition, running speed as a behavioral measure of familiarity (Frank et al.,2004) also significantly increased during training (p < 0.0001, F_(14,154)_ = 11.6). This simple task thus allowed us to investigate experience-dependent changes in hippocampal representations of salient locations by cellular resolution functional imaging.

### Initial establishment of hippocampal salience maps

We next sought to visualize how hippocampal CA1 place maps emerge from a naive state during training on the virtual endless linear track task, with a particular interest in whether and when the representations of the two salient locations become prominent in the map. Two-photon imaging of the CA1 pyramidal cell layer through an optical window in Thy1-G-CaMP7 mice provided an image of a large number of G-CaMP7-labeled pyramidal cells across the entire field of view (Figure 2A). Because the neuronal density of the dorsal CA1 pyramidal cell layer is very high, we computationally extracted the morphology and activity of individual neurons from time-lapse movies using a modified non-negative matrix factorization algorithm to ensure objective and reproducible image analysis (Vogelstein et al., 2010; Pnervmatikakis et al., 2016; Takekawa et al., 2017) (Figure S2). This algorithm assumes that time-varying fluorescence signals of each cell can be decomposed into the product of a spatial filter and a time variation of fluorescence intensity, which are estimated by two alternating iterative steps (for details, see Methods). The spatial filter represents the position and shape of the cell (Figure S2A-B). The timing of spiking activity is inferred using the assumption that each spike evokes transient elevations of fluorescence intensity with a double-exponential shape (Figure S2C).

**Figure 2.**
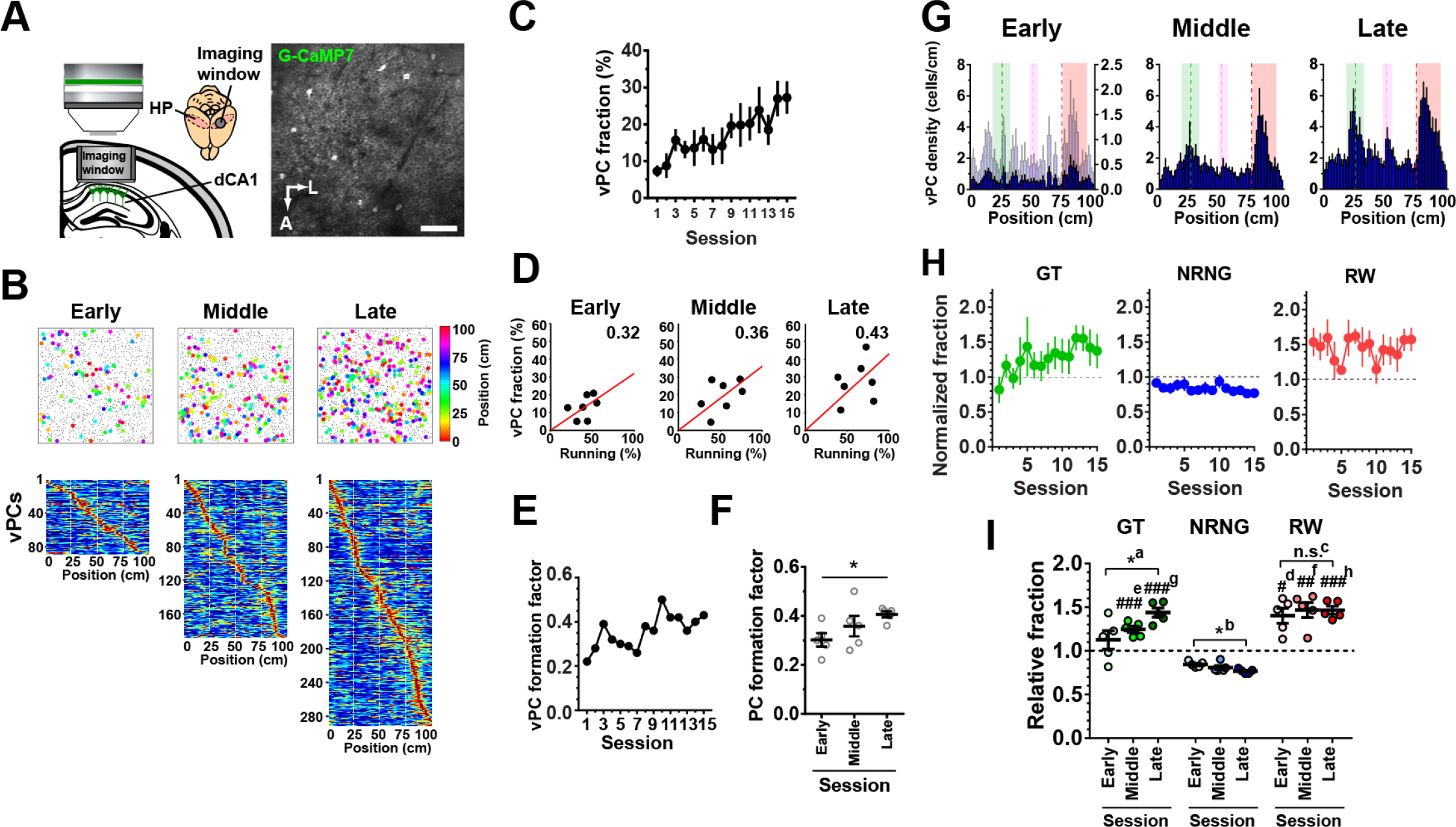
Establishment of salience maps in the hippocampal CA1 deep sublayer. (A) *In vivo* two-photon imaging of G-CaMP7-expressing CA1 pyramidal neurons through a hippocampal imaging window. A stainless cylindrical imaging window with a glass coverslip attached to the bottom was implanted above the dorsal CA1 (dCA1) area of the hippocampus (HP) after aspiration of the overlying cortical tissue (left). A representative fluorescence image of G-CaMP7-expressing pyramidal neurons in the dorsal CA1 hippocampus (right). Active cells are shown as bright cells in this grayscale image. Scale bar = 100 μm. A, anterior;, lateral. (B) Examples of virtual place cell (vPC) maps imaged in the same animal at the early (session 1), middle (session 7) and late (session 14) phases of training on the virtual endless linear track task (top). vPCs and non-vPCs (NvPC) are represented by filled circles of various colors and gray dots, respectively. The different colors of the filled circles represent different locations of the virtual place fields. Heat maps shown below are distributions of virtual place fields of the corresponding sessions ordered by their positions (bottom). (C) The fractions of vPCs relative to the number of total identified cells imaged at each session. (D) Example scatter plots showing the relationship between the fraction of vPCs and the fraction of time spent running for early (session 4), middle (session 9) and late (session 15) phases of the training. The red line in each panel represents linear regression. The value shown at top right indicates the vPC formation factor, which is defined as the slope of the regression line. (E) Changes of vPC formation factors during training. (F) Averages of vPC formation factors for early (sessions 1 - 5), middle (sessions 6 - 10) and late (sessions 11 - 15) phases of training. *P = 0.030, ANOVA with post hoc test for linear trend, n = 5 sessions each. (G) Histograms indicating the distribution of vPCs with respect to the track position for early (session 1), middle (session 6) and late (session 12) phases of the training. The average data from 7 mice are shown. For comparison, the histogram of the early phase was scaled by its maximum value to that of the late phase and is plotted in light blue on its right Y-axis. The green, red and magenta dashed lines delineate the positions of the landmark, reward delivery and boundary of different wall patterns, respectively. The areas shown in green, red and magenta indicate those that define gate, reward and wall cells, respectively. (H) Hippocampal spatial representations as expressed by the fractions of gate cells (GT, green), non-reward, non-gate vPCs (NRNG, blue), and reward cells (RW, red) relative to the number of total vPCs identified in each session. Values were normalized to that obtained in the case of uniform distribution (i.e., 0.0125/bin), and values greater than 1 indicate that the locations are over-represented. (I) Average normalized fractions of GT (green), NRNG (blue) and RW (red) for the early, middle and late phases of the training. *^a^, P = 0.017, F_(2,12)_ = 4.91; *^b^, P = 0.017, F_(2,12)_ = 4.79; n.s.^c^, P = 0.78, F_(2,12)_ = 0.247; one-way ANOVA, n = 5 sessions each; #^d^ P = 0.011 vs. NRNG Early, F_(1.101, 4.405)_ = 8.40; ###^e^, P = 0.0006 vs. NRNG Middle; ##^f^, P = 0.0076 vs. NRNG Middle, F_(1.138, 4.550)_ = 26.5; ###^g^, P = 0.0006 vs. NRNG Late; ###^h^, P = 0.0006 vs. NRNG Late, F_(1449, 5.795)_ = 73.5; one-way ANOVA; n = 5 sessions each.

Using these activity time traces, we determined the virtual location-specific activity by statistical testing of the mutual information content calculated between each cell’s neuronal activity and the animal’s virtual locations (Figure S2D-E). Accumulating evidence has demonstrated that “place cells” in the VR recapitulate many, though not all, place cell properties observed in real environments (Chen et al., 2013; Ravassard et al., 2013). We thus called the cells that exhibited place cell-like virtual location-specific activity “virtual place cells (vPCs)” in this study and used them as a means to understand how information regarding the spatial aspect of the external world is represented in the hippocampal map.

Consistent with previous studies in real and virtual environments, vPCs were formed rapidly within the first session on the virtual linear track (Hill, 1978; Wilson and McNaughton, 1993; Frank et al., 2004; Chen et al., 2013) (Figure 2B-C). The initial fractions of vPCs were low but then increased as the training proceeded. The fraction of vPCs and that of time spent running showed a good overall correlation (Figure S3A). The parallel increase in these two factors during training thus suggested that the observed increase in vPCs might be simply due to more sensitive detection of vPCs, which was enabled by an increase in the length of data for virtual place field calculations. Thus, we created scatter plots of the fraction of vPCs against that of the time spent running for all seven animals for each of the 15 sessions and determined the slopes of the regression lines to obtain indices (termed the vPC formation factor) representing the amount of vPCs formed by a unit length of time spent running (Figure 2D-F). The vPC formation factor significantly increased in the late phase of training compared with the early phase (Figure 2D-F). These results indicate that training indeed facilitated more effective formation of vPCs for a given amount of spatial experience. A separate analysis demonstrating that the sessions in the late phase contained larger fractions of vPCs than the sessions in the early phase with a comparable extent of running time also supports this idea (Figure S3B-C).

We then examined whether the virtual locations associated with salience were disproportionately over-represented in the hippocampal spatial map. The histograms of vPCs against the track position typically exhibited two large peaks, which were clearer in the late training phase; one peak corresponded to the location of the landmark (i.e., the green gate), and the other corresponded to that of the reward (Figure 2G). While the first peak closely matched the landmark location, the second peak was slightly shifted to the direction of the mouse’s running, which likely reflects that the animals recognized or anticipated the rewards at places that were slightly past the delivery point, as suggested by slowing down of running speed around this zone (data not shown). Importantly, the over-representation of the location for the reward was discernible even in the first session of training, whereas that of the location for the landmark gradually developed as the training proceeded (Figure 2G-I). The fraction of vPCs that encoded the location of the reward (here termed “reward cells” (RW) for convenience) was not significantly different between the early and late phases of training, whereas that of vPCs that encoded the location of the landmark (similarly termed “gate cells” (GT)) increased significantly with a complementary decrease in the fraction of vPCs that encoded other locations (termed “non-reward, non-gate vPCs” (NRNG)) (Figure 2H-I). The slower increase in vPCs that encode locations associated with salient visual cues is further supported by a more delayed and smaller increase in vPCs that encode a location with less visual salience, such as a boundary of different wall patterns (termed “wall cells” (WL), Figure S4). Collectively, these results demonstrate that the over-representation of salient locations is formed and maintained at a population level, even though the maps develop dynamically throughout the training period. The establishment and refinement of representations of salience depend on its nature; the representation of motivational salience is established rapidly, whereas that of environmental salience develops over the course of training.

What is the benefit of this salience representation in encoding and storage of information about the animal’s environment? We trained a Bayesian decoder from the first 150 s of the running period and reconstructed the animal’s trajectory in the following 90 s using the activities of vPCs in the same sessions (Zhang et al., 1998; Ziv et al., 2013) (Figure 3A). The average median errors across all track positions were initially large in the early phase of training and decreased steadily as the training proceeded (Figure 3B), likely due to the training-induced increase in vPC number (Figure 3C). A more detailed analysis of well-decoded sessions (average median error across all positions < 10 cm) revealed that the errors for the locations associated with the landmark and reward were significantly smaller than those for other locations (Figure 3D). These results demonstrate that the established salience maps enable more accurate population coding for these locations.

**Figure 3.**
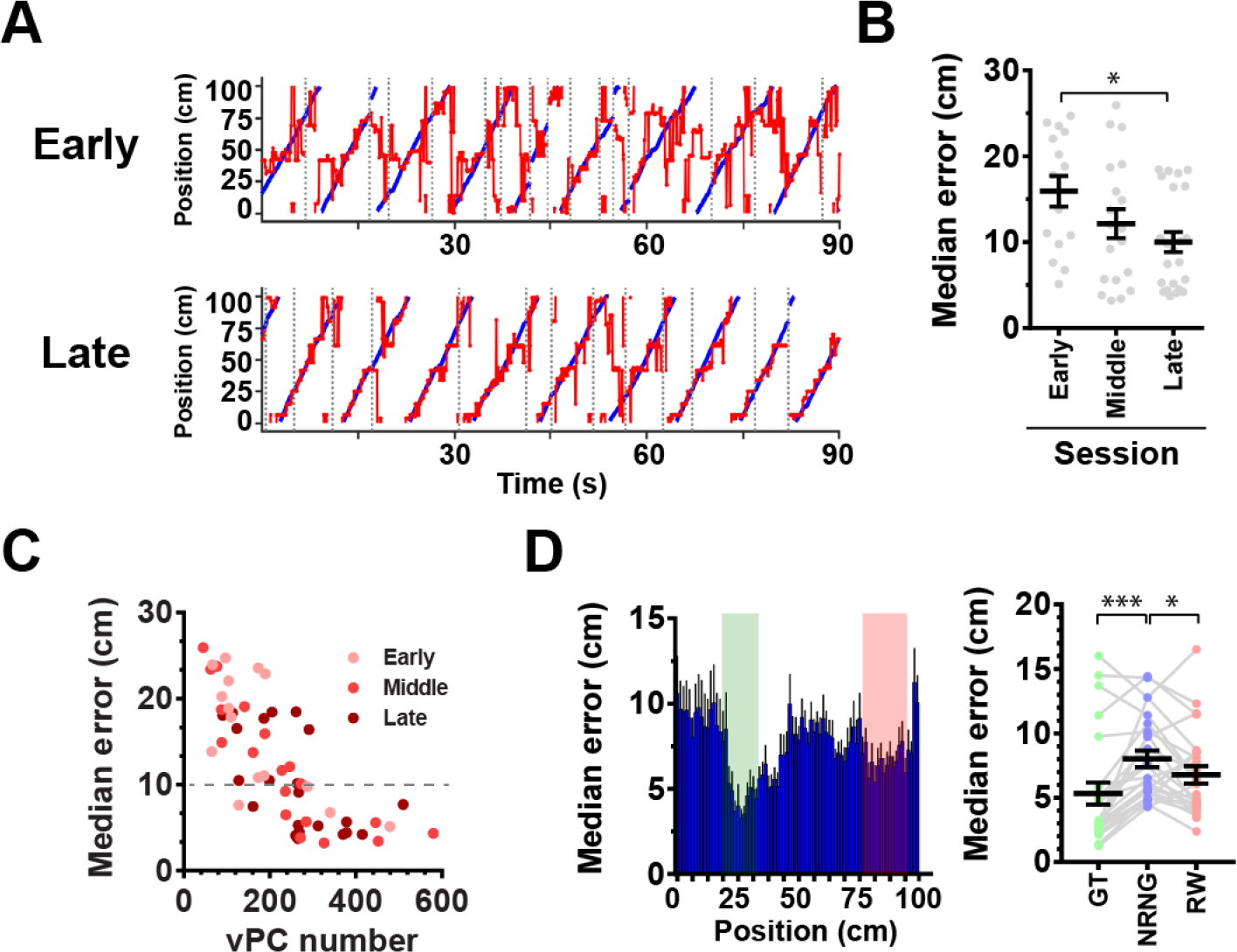
Precise estimation of salient locations by over-represented CA1 salience maps. (A). Examples of trajectories estimated by the Bayesian decoder for early (top) and late (bottom) phases of training. The results of the entire 90-second test periods from the same animal are shown. Blue and red lines represent real and estimated trajectory, respectively. Periods of immobility that separated continuous running were excluded from decoding and their positions are indicated by vertical dashed lines. (B) Average median errors for the early, middle and late phases of training. *P = 0.018, F_(2,54)_ = 3.65, one-way ANOVA, n= 15, 19 and 23 sessions. (C). Relationship between absolute vPC numbers and average median estimation errors across the locations. The symbols filled with pink, red and dark red indicate sessions in the early, middle and late phases, respectively. The dashed line represents the threshold for well-decoded sessions. Only the sessions with running times ≧240 s were analyzed (n= 57 sessions). (D) Average median errors across all well-decoded sessions plotted against the position (left) and those for the locations encoded by GT, NRNG, and RW (right). *P = 0.026, ***P< 0.0001, F_(1.312, 31.49)_ = 8.14; one-way ANOVA, n = 25 sessions.

### Experience-dependent map consolidation

The vPC maps imaged at each session appeared rather different from each other, even within the same animals, implying that hippocampal spatial representations are highly dynamic while being established (Figure 2B). This result further suggests that while new representations were created by each experience, at least in parts of the maps, preexisting representations are either eliminated or stabilized. To gain insight into whether repeated training stabilizes the maps, we next investigated training-induced changes in the maps at an individual cell level by comparing the virtual place fields of the same cells across different sessions (Figure S5). In this analysis, we conservatively focused on comparisons between two consecutive sessions (i.e., sessions 1 and 2, 2 and 3, etc.) because the quality of the image alignment was gradually reduced as the number of sessions that separated the two images increased (P < 0.0001, F_(13, 721)_ = 10.94, one-way ANOVA, Figure S5F). In the early maps, which contained a relatively small fraction of vPCs, only a small number of vPCs were identified as common in both sessions (hereafter called “common vPCs”), whereas the fraction of common vPCs increased significantly as more vPCs were imaged in the late phase of the training (Figure 4A-C). Moreover, the fraction of vPCs that had stable virtual place fields (< 10 cm difference) in both sessions (“stable vPCs”) also increased markedly as the training proceeded (Figure 4A-B, D-E), indicative of experience-dependent map consolidation. Image comparisons between adjacent sessions showed that fractions of common cells (i.e., cells that were identified in common in the two sessions) were constant over time (P = 0.57, repeated measures one-way ANOVA) (Figure S5E), verifying that the increased stability of vPCs was not caused by differences in image alignment. Furthermore, the fractions of common and stable vPCs normalized by the number of vPCs also increased significantly as the training proceeded, revealing that the training-induced increase of vPC stability was not simply due to the increase of the number of vPCs (Figure S6). We then asked whether representations of salient locations are more stable than those of non-salient locations. We calculated the fractions of stable vPCs with respect to the number of common vPCs as an index for stable representations at each location and found that this index was significantly higher for locations associated with the landmark or reward than for other locations (Figure 4F-G). Finally, we tested the hypothesis that the stability of the maps predicts the performance of animals in the behavioral task. We found that the gain of vPC stability between the early and late phases of the training in individual animals exhibited a good linear correlation with their differences in time spent running between the two phases of the training (r = 0.84, Figure 4H). The results suggest that hippocampal place maps are more strongly stabilized if the animals more effectively learn to run the virtual linear track task.

**Figure 4.**
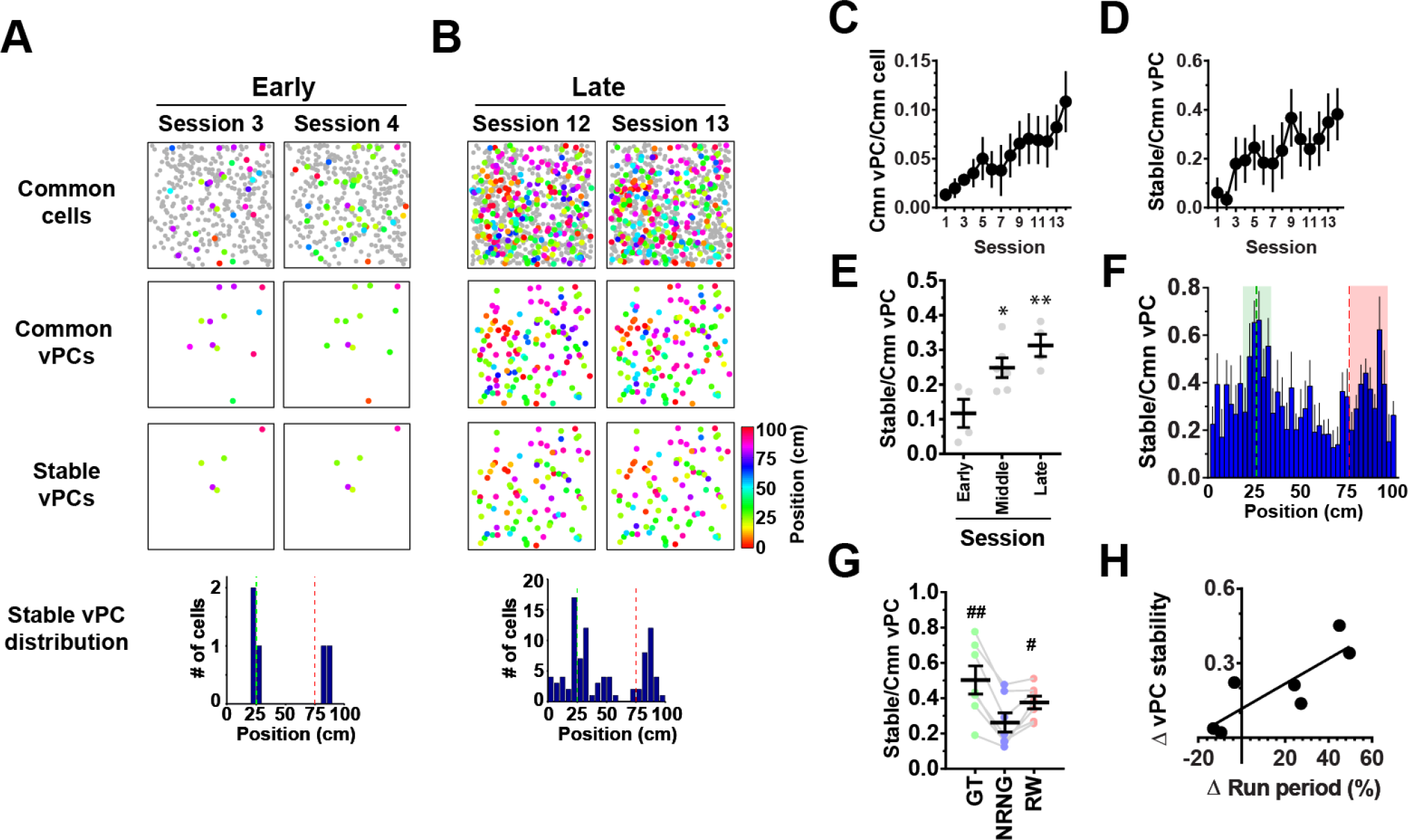
Experience-dependent consolidation of salience maps. (A) Example hippocampal CA1 vPC maps imaged in two consecutive sessions in the early phase of the training. Maps shown on top, middle and bottom present cells identified in common in both sessions (Common cells), cells identified as vPCs in both sessions (Common vPCs), and cells identified as vPCs with stable (< 10 cm difference) virtual place fields in both sessions (Stable vPCs), respectively. vPCs and non-vPCs are represented by filled circles of various colors and gray dots, respectively. The different colors of the filled circles represent different locations of the virtual place fields. The histogram shown at the bottom indicates the distributions of the stable vPCs against the track position. The green and red dashed lines delineate the positions of the landmark and reward delivery, respectively. The same convention applies to B. (B) vPC maps imaged in the late phase of training in the same animal as presented in a. (C) The fraction of common vPCs relative to the number of common cells identified in the two consecutive sessions that were compared. The X-axis indicates the earlier of the two sessions that were compared. (D) vPC stability calculated as the fraction of stable vPCs relative to the number of common vPCs identified in the two consecutive sessions that were compared. (E) Average vPC stability for the early (sessions 1 - 4, which indicates the earlier of the two sessions that were compared), middle (sessions 5 - 10) and late (sessions 11 - 14) phases of training. *P = 0.029 vs. Early, **P = 0.0048 vs. Early, F_(2,11)_ = 7.90, one-way ANOVA; n = 4, 6 and 4 session pairs. (F) The average fractions of stable vPCs relative to the number of common vPCs plotted against the track position. Values were calculated from data across all sessions and averaged for 7 mice. The green and red dashed lines delineate the positions of the landmark and reward delivery, respectively. The areas shown in green and red indicate those that define gate and reward cells, respectively. (G) The average vPC stability for gate cells (GT), non-reward, non-gate vPCs (NRNG) and reward cells (RW). ^#^P = 0.029 vs. NRNG, ^##^P = 0.0018 vs. NRNG, F_(1.202, 7.212)_ = 13.9, one-way ANOVA, n = 7 mice from 2 groups. (H) The relationship between vPC stability and task performance. The X-axis presents the task performance of each mouse measured by the difference in the fraction of time spent running between the early (average of sessions 1 - 5) and late (average of sessions 11 - 15) phases of the training. The Y-axis presents the difference in vPC stability between the early and late phases of training.

### Cellular dynamics for map establishment

To elucidate the cellular mechanism for the map establishment, we conducted two analyses of the functional transitions of cells belonging to different virtual location-related cell categories between sessions (Figures 5 and S7). In the first analysis, common cells in a reference session N were classified into vPCs or non-vPCs (NvPC) according to their virtual location-related activities, and the probabilities of transitioning to the same or the other categories in the subsequent session N+1 were calculated (for details, see Methods) (Figure S7A). The probability of vPC to remain as a vPC, *P* _vpc-vpc_, and that of “*de novo*“ vPC formation from NvPC, *P* _NvPC-vPC_, increased significantly between the early and late phases of training (Figure S7B-C), with complementary decreases in the probabilities of vPC disappearance *P* _vpc-Nvpc_ and NvPC stabilization *P* _Nvpc-Nvpc_ (Figure S7B). In addition, *P* _vPC-vPC_ was significantly greater than *P* _NvPC-vPC_ in both training phases (Figure S7B-C). This result implies that vPCs are more likely to be vPCs in the subsequent sessions than NvPCs are, and that such a stabilization process plays a major role in the increase in vPCs. Together, these results demonstrate that training induces expansion of the vPC population by shifting the balance of a dynamic exchange between vPCs and NvPCs toward the direction of vPC stabilization and formation.

**Figure 5.**
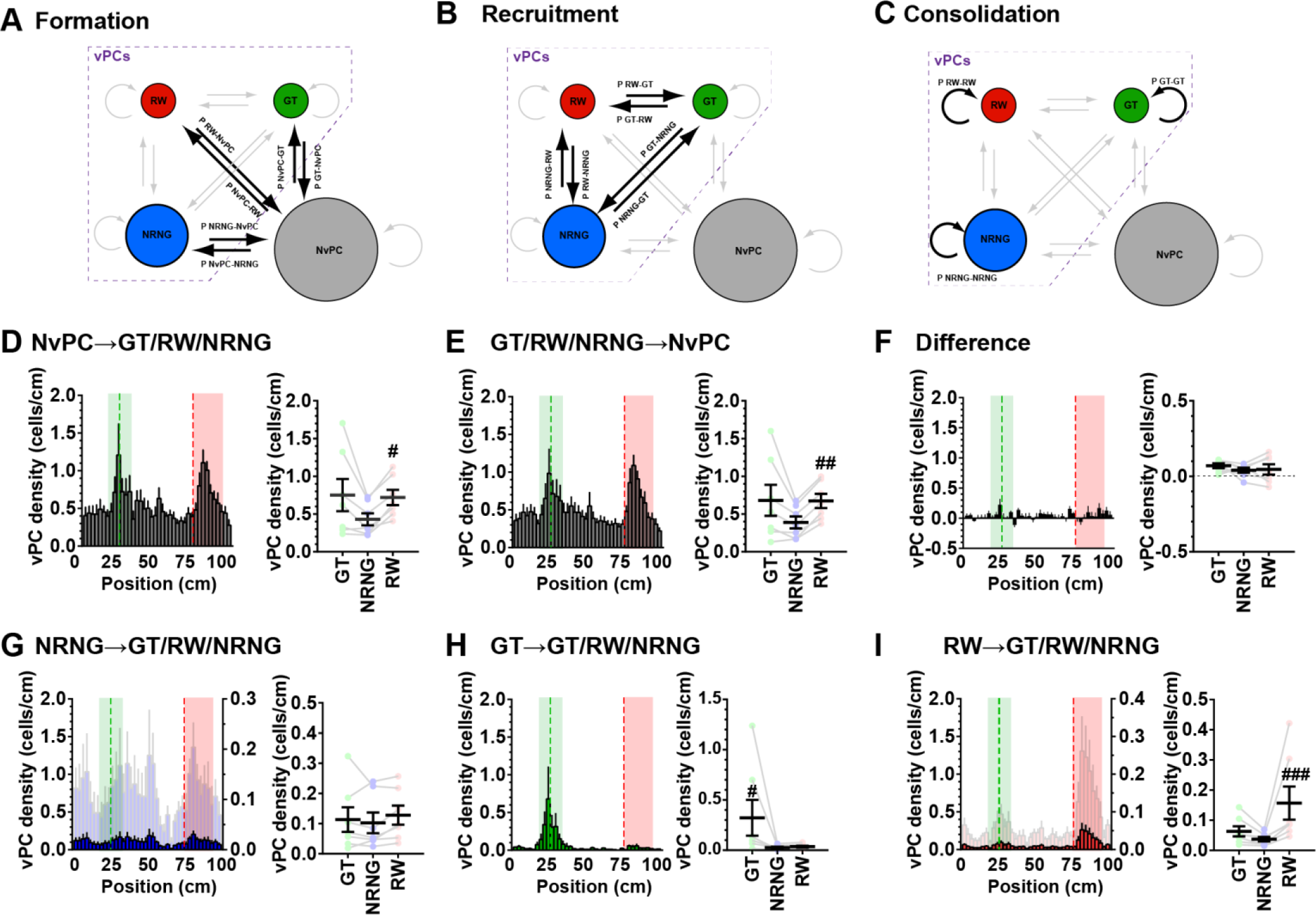
Formation, recruitment and stabilization of different vPC categories during map establishment. (A-C) Schematic diagrams of direct formation (A), lateral recruitment (B) and selective consolidation models (C). In each diagram, four functional cell categories, namely, reward cells (RWs, red), gate cells (GTs, green), non-reward, nongate vPCs (NRNGs, blue) and non-vPCs (NvPCs, gray), and the transitions between them are defined. RWs, GTs and NRNGs constitute subcategories of vPCs and are shown enclosed by a dashed line. The transitions relevant to each model are highlighted in black. (D) Formation of different vPC categories from NvPCs. (left) A histogram showing the distribution of vPCs that were NvPCs in the previous sessions against the track position. The values were calculated from data across all sessions and averaged for 7 mice. For comparison, the histograms shown in D, E and G-I are plotted on the left Y-axes on the same scale. In addition, the histograms in G and I were scaled by their maximum values and plotted in a light color on the right Y-axes. (right) The average cell density of each vPC subcategory formed from NvPCs. ^#^P = 0.023 vs. NRNG, χ^2^_(2)_ = 8.00; Friedman test, n= 7 mice from 2 groups. (E) Elimination of different vPC categories into NvPCs. (left) A histogram showing the distribution of vPCs that became NvPCs in subsequent sessions against the track position. (right) The average cell density of each vPC subcategory that became NvPCs in the subsequent sessions. ^##^P = 0.0099 vs. NRNG, χ^2^_(2)_ = 8.86; Friedman test, n = 7 mice from 2 groups. (F) Net vPC formation. (left) A histogram of the difference obtained by subtracting the histogram in E from that in D. Note that this histogram only shows the distribution of locations for entry into and exit from the vPC populations and can have negative values for some locations. (right) The average cell density of the remaining vPCs classified by the vPC subcategories. (G) Transition and stability of the NRNGs. (left) A histogram showing the distribution of vPCs that were NRNGs in the previous sessions against the track position. (right) The average cell density of each vPC subcategory that was derived from NRNG. (H) Transition and stability of the GTs. (left) A histogram showing the distribution of vPCs that were GTs in the previous sessions against the track position. (right) The average cell density of each vPC subcategory that was derived from GTs. ^#^P = 0.023 vs. NRNG, χ^2^_(2)_ = 7.14, Friedman test, n = 7 mice from 2 groups. (I) Transition and stability of the RWs. (left) A histogram showing the distribution of vPCs that were RWs in the previous sessions against the track position. (right) The average cell density of each vPC subcategory that was derived from RWs. ^###^P = 0.0005 vs. NRNG, χ^2^_(2)_ = 14.0, Friedman test, n = 7 mice from 2 groups.

Next, to clarify the contributions of the formation, recruitment and consolidation of vPCs in the establishment of salience maps (Figures 1A-C and 5A-C, see also Introduction), we further subclassified the vPC population into RW, GT and NRNG according to their virtual place field positions and analyzed the transitions between them for each vPC subcategory in the second analysis. The formation of vPCs from NvPCs exhibited significant biases toward RWs (Figure 5D). However, a subpopulation of vPCs that became NvPCs in subsequent sessions also exhibited a similarly biased distribution (Figure 5E). The net effect, calculated as their difference, resulted in no significant biases (Figure 5F), implying that the disproportionate formation of vPCs predicted by the “direct formation model” (Figure 1A) exists but is counteracted by a similar bias in the disappearance of part of the vPCs. In addition, the distribution of vPCs derived from former NRNGs appeared not to be biased toward RWs or GTs but was rather uniform (Figure 5G), providing evidence against the “lateral recruitment model” (Figure 1B). In contrast, the distribution of vPCs derived from former GTs or RWs was substantially biased toward the location by which each of the two vPC subcategories was defined, as predicted by the “selective consolidation model” (Figures 1C and 5H-I). These results further confirm that RWs and GTs have a marked tendency to persistently encode the same locations and strongly support the idea that selective consolidation of relevant spatial representations is the primary mechanism underlying the establishment of hippocampal salience maps.

### Robust map reorganization by conjunction of salience

Our findings for experience-dependent map establishment thus far imply that the stability of spatial representations in the hippocampal map can be controlled by the presence of salient features. This result raises a further question concerning how this process is engaged in the plasticity of pre-established maps, which was addressed by imaging the salience maps during re-training in the reward rearrangement task. In this task, mice trained on the standard virtual linear track (i.e., the track on which visual landmark and reward delivery were associated with two distinct locations) for 15 sessions were further trained for the following 5 sessions in the same linear track, except that the location of reward delivery was shifted to the landmark location (Figure 6A). This task allowed us to examine the changes in the map that occurred when mice re-experienced the environment in which two separately located salient features were now jointly presented. We divided the entire re-training period into early (session 1-2) and late (session 3-5) sessions. Distance traveled, running speed and fraction of time spent running were not significantly changed between before and after the reward rearrangement (Figure S8A-B). However, this manipulation triggered robust reorganization of the salience maps (Figure 6B). Notably, the fraction of vPCs decreased immediately after reward rearrangement but recovered as the re-training proceeded (Figures 6B and S8C). Furthermore, the over-representation of the previous reward location disappeared suddenly as early as the first rearrangement sessions, whereas over-representation of the location associated with the conjunction of the pre-existing landmark and the relocated reward was markedly enhanced over the course of re-training (Figure 6C-E). The effect of this enhanced over-representation appeared rather additive (normalized fractions of vPCs, Pre GT, 1.14 ± 0.10; Pre RW, 1.65 ± 0.09; Late GT+RW, 1.89 ± 0.18), supporting the view that the magnitude of increases in place cell numbers represents the degree of salience in the hippocampus map. Interestingly, the reward rearrangement also enhanced the representation of the location associated with the wall pattern transition (Figure 6C-E), demonstrating that a salience conjunction can also be formed broadly with a nearby less-salient visual cue.

**Figure 6.**
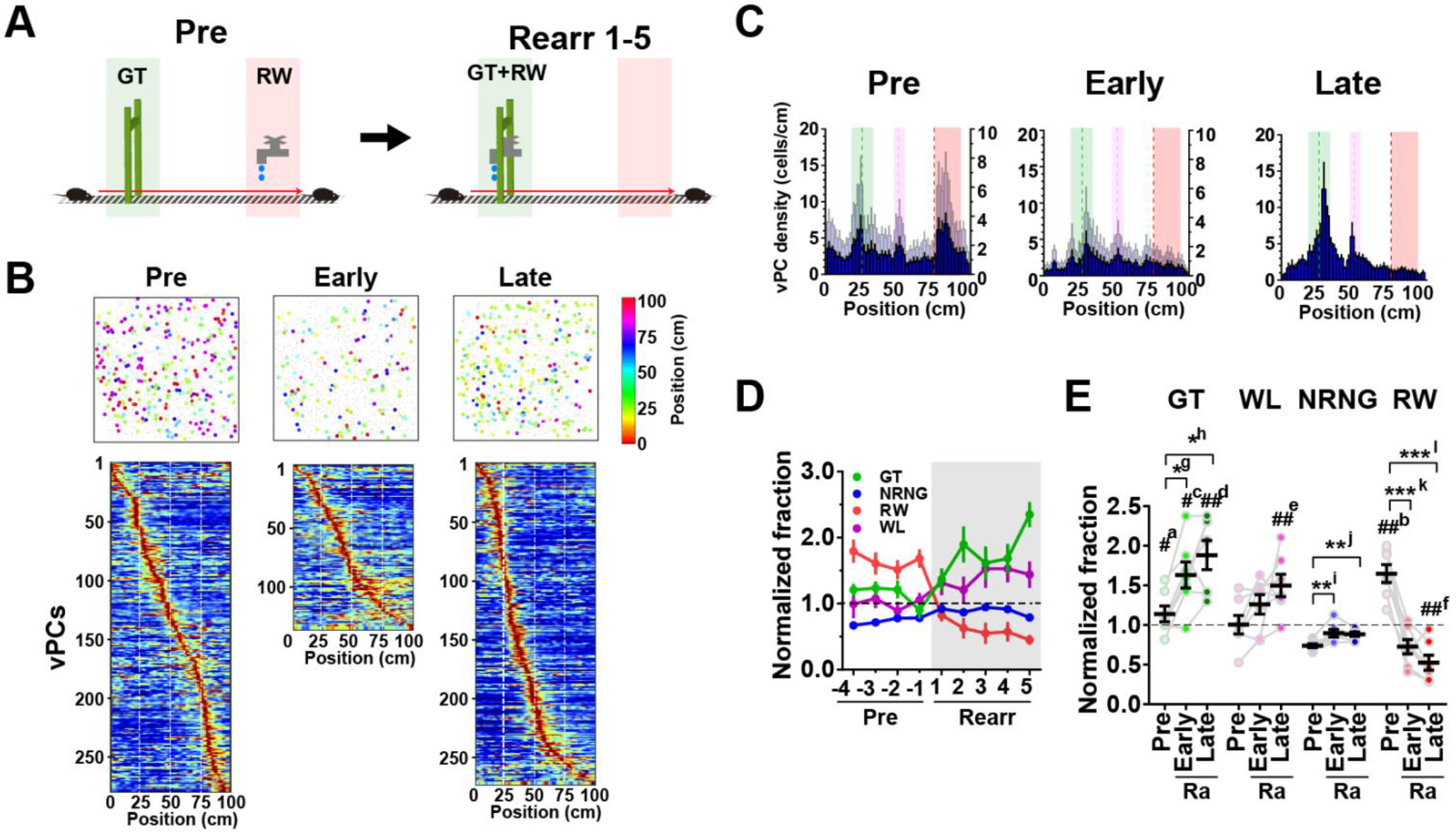
Robust reorganization of salience maps induced by a conjunction of different kinds of salience. (A) Design of the reward rearrangement task. Mice were first trained in the standard linear track that included a visual landmark (GT) and reward delivery (RW) at separate locations for 15 sessions (Pre, left). Once training was complete, the location of reward delivery was shifted to match the location of the visual landmark (GT+RW), and the mice were re-trained in this new arrangement for the following 5 sessions (Rearr 1-5, right). (B) Examples of vPC maps imaged in the same animal in the pre (session Pre -1), early (session Rearr 1) and late (session Rearr 5) phases of the reward rearrangement task (top). vPCs and non-vPCs (NvPC) are represented by filled circles of various colors and gray dots, respectively. The different colors of the filled circles represent different locations of the virtual place fields. Heat maps shown below the vPC maps are the distributions of virtual place fields of the corresponding sessions ordered by their positions (bottom). (C) Histograms indicating the distribution of vPCs with respect to the track position for pre (session Pre -2), early (session Rearr 1) and late (session Rearr 5) phases of the rearrangement task. The average data from 7 mice are shown. For comparison, the histograms of the pre and early sessions were scaled by their maximum values to that of the late session and are plotted in light blue on their right Y-axes. The green, red and magenta dashed lines delineate the positions of the landmark, reward delivery and boundary of different wall patterns, respectively. The areas shown in green, red and magenta indicate those that define gate, reward and wall cells, respectively. (D) Hippocampal spatial representations as expressed by the fractions of gate cells (GT, green), non-reward, non-gate vPCs (NRNG, blue), reward cells (RW, red) and wall cells (WL, magenta) relative to the number of total vPCs identified in each session. Values were normalized to that obtained in the case of uniform distribution (i.e., 0.0125/bin), and values greater than 1 indicate that the locations are over-represented. (E) Average normalized fractions of GT (green), WL (magenta), NRNG (blue) and RW (red) for pre, early and late phases of the reward rearrangement (Ra) task. #^a^, P = 0.044 vs. NRNG Pre, ##^b^, P = 0.0022 vs. NRNG Pre, F_(1.433, 8.597)_ = 13.4; #^c^, P = 0.026 vs. NRNG Early, F_(2.019, 12.11)_ = 10.8; ##^d^, P = 0.0087 vs. NRNG Late, ##^e^, P = 0.0095 vs. NRNG Late, ##^f^, P = 0.0095 vs. NRNG Late, F_(1.785, 10.71)_ = 18.3; *g, P = 0.011, *h, P = 0.011, F_(1.511, 9.067)_ = 10.5; **i, P = 0.0099, **j, P = 0.0041, F_(1.672, 10.03)_ = 11.6; ***k, P = 0.0008, ***l, P = 0.0008, F_(1.408,8.446)_ = 38.8; one-way ANOVA, n= 7 mice from 2 groups.

Cell-by-cell comparisons of the maps imaged at adjacent sessions further revealed salience-dependent, location-specific regulation of hippocampal map stability (Figure 7A-D). The reward rearrangement triggered a significant reduction in stability for the representation of the previous reward location and the non-salient locations in the early phase of re-training, while the stability of the location with conjunctive salience was essentially maintained (Figure 7B and D). The stability of the non-salient locations recovered in the late phase, but that for the previous reward location remained low (Figure 7C-D). These results demonstrate that the hippocampal map plasticity accompanies a redistribution of vPC stability through a short period of heightened map instability. The presence and absence of salience thus govern the hippocampal map representations through a dynamic regulation of place field stability.

**Figure 7.**
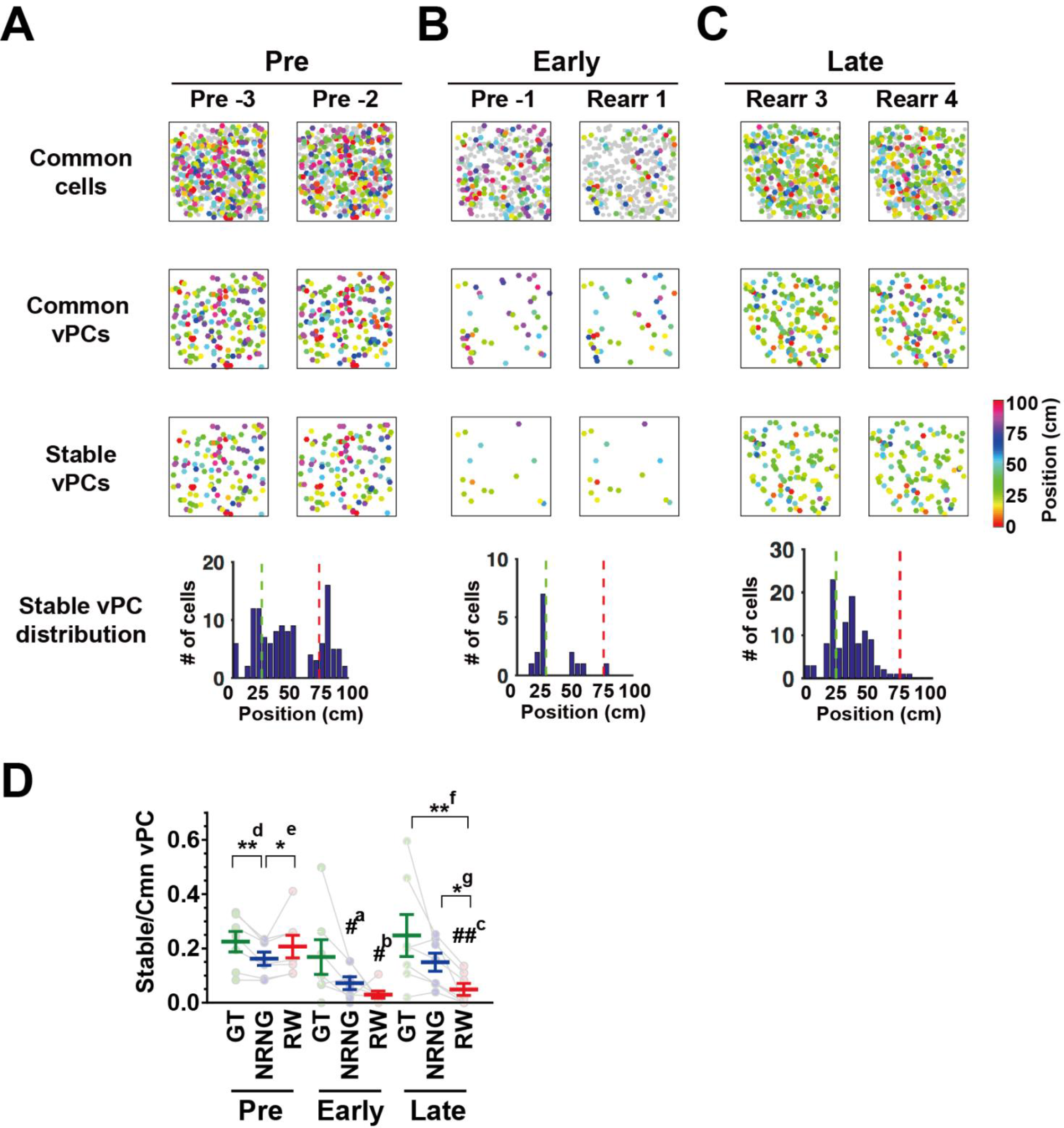
vPC stability during reward rearrangement task. (A) Example hippocampal CA1 vPC maps imaged in two consecutive sessions in the pre phase of the reward rearrangement task. Maps shown on top, middle and bottom present cells identified in common in both sessions (Common cells), cells identified as vPCs in both sessions (Common vPCs), and cells identified as vPCs with stable (< 10 cm difference) virtual place fields in both sessions (Stable vPCs), respectively. vPCs and non-vPCs are represented by filled circles of various colors and gray dots, respectively. The different colors of the filled circles represent different locations of the virtual place fields. The histogram shown at the bottom indicates the distributions of the stable vPCs against the track position. The green and red dashed lines delineate the positions of the landmark and reward delivery, respectively. The same convention applies to B and C. (B-C) Hippocampal CA1 vPC maps imaged in the early (B) and late (C) phase of training in the same animal as presented in A. (D) The average vPC stability for GT, NRNG and RW for pre, early and late phases of the reward rearrangement task. #^a^, P = 0.049 vs. NRNG Pre, χ^2^_(2)_ = 6.00; #^b^, P = 0.033 vs. RW Pre, ##^c^, P = 0.0063 vs. RW Pre, χ^2^_(2)_ = 11.2; **d, P = 0.0040, *e, P = 0.049, χ^2^_(2)_ = 11.1; **f, P = 0.0040, *g, P = 0.049, χ^2^_(2)_ = 11.1; Friedman test, n = 7 mice from 2 groups.

### Cellular dynamics for map reorganization

Finally, to elucidate the cellular mechanism underlying the salience rearrangement-induced map plasticity, we analyzed the contribution of direct formation, lateral recruitment and selective consolidation of vPCs during this process (Figures 1A-C and 8). For simplicity of terminology, cell categories are labeled based on the condition before reward rearrangement. More specifically, GT and RW cells refer to the vPCs that encode the landmark and reward locations at the time before reward rearrangement, and their field positions maintain the same designations so that they become associated with the landmark plus reward and no salience after reward rearrangement, respectively. The reward rearrangement elicited a rapid and sustained reduction in *de novo* formation of RW from NvPCs in the early and late phases and a delayed increase in *de novo* GT formation in the late phase of re-training (Figure 8A-i and B-i). The elimination of RW into NvPCs subsided in the late phase (Figure 8A-ii and B-ii), likely reflecting the substantial reduction of the RW formation in the preceding early phase. Consequently, the net effect gave rise to a remarkable transient decrease in RW formation in the early phase and a delayed increase in GT formation (Figure 8A-iii and B-iii). These results demonstrate that the reward rearrangement and the formation of conjunctive salience markedly influence the net vPC formation and that this salience-dependent down-and up-regulation of vPC formation underlies the rapid decrease in the vPC fraction and disappearance of the overrepresentation of the previous RW location in the early phase and the recovery of the vPC fraction and enhancement of the over-representation of the GT location in the late phase of plasticity.

**Figure 8.**
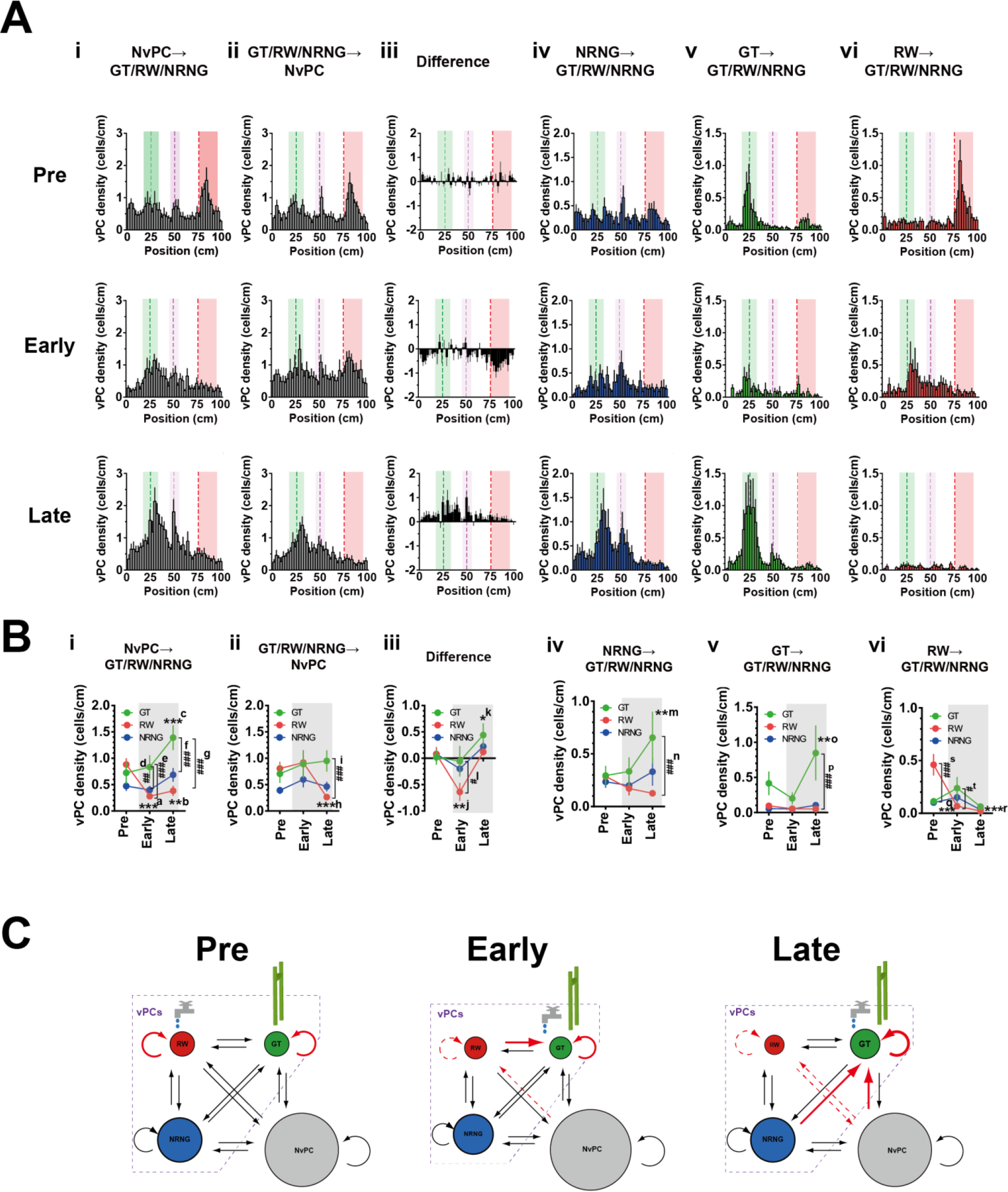
Map plasticity is mediated by a dynamic interplay of stabilization, formation and recruitment of vPCs. (A-i) Formation of different vPC categories from NvPCs. The histograms show the distributions of vPCs that were NvPCs in the previous sessions against the track position, and those for pre, early and late phases of the reward rearrangement task are shown from top to bottom. The values were calculated from data across all relevant sessions and averaged for 7 mice. The green, red and magenta dashed lines delineate the positions of the landmark, reward delivery and boundary of different wall patterns, respectively. The areas shown in green, red and magenta indicate those that define gate, reward and mid cells, respectively. (A-ii) Elimination of different vPC categories into NvPCs. The histograms show the distributions of vPCs that became NvPCs in the subsequent sessions against the track position. (A-iii) Net vPC formation. The histograms of the differences were obtained by subtracting the histograms in A-ii from the corresponding ones in A-i. (A-iv) Transition and stability of the NRNGs. The histograms show the distributions of vPCs that were NRNGs in the previous sessions against the track position. (A-v) Transition and stability of the GTs. The histograms show the distributions of vPCs that were GTs in the previous sessions against the track position. (A-vi) Transition and stability of the RWs. The histograms show the distributions of vPCs that were RWs in the previous sessions against the track position. (B-i) Formation of different vPC categories from NvPCs. The average cell density of each vPC subcategory formed from NvPCs are shown for pre, early and late phases of the reward rearrangement task. The periods of reward rearragement are shaded in gray. ***a, P = 0.0002 vs. RW Pre, **b, P = 0.0015 vs. RW Pre, ***c, P < 0.0001 vs. GT Pre, ##d, P = 0.0055, ###e, P = 0.0005, ###f, P < 0.0001, ###g, P < 0.0001, F_(2,12)_ = 14.5, two-way ANOVA, n = 7 mice from 2 groups. (B-ii) Elimination of different vPC categories into NvPCs. ***h, P = 0.0007 vs. RW Pre, ###i, P < 0.0001, F_(2,12)_ = 8.01, two-way ANOVA, n = 7 mice from 2 groups. (B-iii) Net vPC formation. The difference between the formation and elimination of each vPC subcategory are shown. **j, P = 0.0019 vs. RW Pre, *k, P = 0.032 vs. GT Early, #l, P = 0.011, F_(2,12)_ = 16.4, two-way ANOVA, n = 7 mice from 2 groups. (B-iv) Transition and stability of the NRNGs. **m, P = 0.0029 vs GT early, n, P < 0.0001, F_(2,12)_ = 5.22, two-way ANOVA, n = 7 mice from 2 groups. (B-v) Transition and stability of the GTs. **o, P = 0.0042 vs. GT Early, ###p, P = 0.0005, F_(2,12)_ = 7.18, two-way ANOVA, n = 7 mice from 2 groups. (B-vi) Transition and stability of the RWs.***q, P < 0.0001 vs. RW Pre, ***r, P < 0.0001 vs. RW Pre, ###s, P < 0.0001, #t, P = 0.036, F_(2,12)_ = 4.71, two-way ANOVA, n = 7 mice from 2 groups. (C) A model for reorganization of the hippocampal salience map elicited by the rearrangement of reward. The key processes in each phase are shown in red. Augmentation and reduction of the processes are shown with thicker and dashed lines, respectively. Selective consolidation of GT and RW plays a dominant role in establishment and maintenance of the salience map during training before the rearrangement (Pre, left). Immediately after reward arrangement, the fraction of vPCs decreases because of decreased RW formation and stabilization, and a subpopulation of RW move their fields to the new reward location (Early, middle). After a few sessions, the stability and formation of GT as well as recruitment from NRNG to GT increase (Late, right).

The major mechanistic differences between map establishment and reorganization are not limited to the regulation of vPC formation. The reward rearrangement significantly increased recruitment of GT from NRNGs toward the late phase of re-training (Figure 8A-iv and B-iv). Furthermore, the recruitment of GT from RW also exhibited a significant transient increase in the early phase of re-training (Figure 8A-vi and B-vi). These results indicate that a conjunction of salience formed by reward rearrangement augments the recruitment of vPCs that encode a previously salient location as well as non-salient locations. The density of stable RW cells exhibited a sudden and prolonged drop in the early and late phases (Figure 8A-vi and B-vi), whereas that of GT remained high throughout the re-training period (Figure 8A-v and B-v), consistent with earlier findings (Figure 7). In summary, the salience rearrangement-induced map reorganization is mediated by a rapid disappearance of the over-representation of the previously salient location followed by a gradual enhancement of the over-representation of the location associated with a newly formed strong conjunction of salience. In contrast to the primary role of selective place field stabilization in initial map establishment, the adaptive change of the hippocampal salience map is cooperatively achieved by multiple forms of cellular dynamics that involve not only stabilization but also direct formation and lateral recruitment of place cells (Figure 8C, see Discussion for details). Notably, GT cells immediately after reward rearrangement were derived more from RW cells than from NRNG cells (46 RW-to-GT cells and 36 NRNG-to-GT cells of 107 total GT cells from 7 mice compared with 22 and 70 cells, which were expected from the uniform distribution across positions, P < 0.0001, Chi-square test). This suggests that RW cells form a unique subpopulation in CA1 and may encode information associated with the reward itself (Figure S9).

## Discussion

Theories of hippocampus-dependent memory and navigation are based on the premise that spatial and non-spatial information are jointly represented in the hippocampus (Knierim et al., 2006; Eichenbaum and Cohen, 2014). Accumulating evidence demonstrates that reward or geometric cues associated with particular locations are represented by an increased density of relevant place cells in hippocampal neural maps, particularly those in the CA1 deep sublayer (O’Keefe and Conway, 1978; Wiener et al., 1989; Hetherington and Shapiro, 1997; Hollup et al., 2001; Dombeck et al., 2010; Dupret et al., 2010; Danielson et al., 2016). By tracking the activity of individual neurons in G-CaMP7 transgenic mice trained in a spatial task in virtual reality, we investigated the cellular principles underlying the formation and plasticity of these maps. We observed that over-representation of salient locations occurred with different time courses and degrees depending on the nature and extent of salience, which resulted in more precise population coding for these locations. We also showed that hippocampal maps were consolidated by experience-dependent stabilization that was correlated with the behavioral performance of the animals. Moreover, stabilization of spatial representations for salient locations but not *de novo* formation and lateral recruitment supported the establishment of the overrepresented maps, providing evidence that place field stability is fine-tuned by experience and salience. Finally, we revealed that reorganization of pre-formed maps was mediated by a coordinated interplay of *de novo* formation, lateral shifts and selective consolidation. These findings provide a comprehensive framework for cellular mechanisms underlying the formation and plasticity of hippocampal functional maps and explain how salient experience can form lasting yet adaptable memory traces.

### Cellular mechanisms for formation and plasticity of hippocampal maps

The observed contrasting cellular dynamics between map formation and plasticity suggests that hippocampal circuits transit between two distinct modes during different experiences. Based on our observations, we propose the following model — during initial map formation, *de novo* place cell formation biased toward salient locations is counteracted by similarly biased place cell removal, whereas over-representations are developed and maintained by a selective consolidation and elevated equilibrium state of salient place cells with little input from neutral place cells by lateral recruitment. This “encoding” mode is dynamically shifted to “updating” mode upon rearrangement of salient features. Thus, the balanced formation and disappearance of place cells is transiently disequilibrated during the early phase of map plasticity to achieve a rapid shift in the peak place cell density. In parallel, a subset of cells that encoded previously salient locations rapidly move their fields to locations with relocated features during the early phase, followed by further recruitment of neutral place cells during the late phase. These processes are accompanied by transient weakening of place cell stability during the early phase and its recovery during the late phase. Thus, our findings indicate that a simple stabilization principle, which allows for encoding salient locations as more persistent traces and allocating more cellular resources to them, governs map formation, whereas parallel and coordinated engagement of multiple processes, which supports the rapid updating of stored information, controls the adaptive reorganization of pre-established maps.

The model assumes that neural signals conveying information about the presence of salience modulate the above processes. For example, salience signals may modulate place cell stability via a dopamine-dependent mechanism (Kentros et al., 2004) and place field formation and disappearance at CA1 pyramidal cell dendrites and EC3 inputs (Bittner et al., 2015; Sheffield et al., 2015). The salience-chasing property observed in this study (i.e., from RW to RW+GT) may arise within or outside the hippocampus (Weible et al., 2009; Deshmukh and Knierim, 2011; Tsao et al., 2013). Remarkably, the lateral recruitment of neutral place cells identified in the late phase of plasticity may be mediated by a mechanism different from salience chasing that likely reflects the binding of cells to a reference frame of salience (Gothard et al., 1996) and may require stronger signals derived from salience conjunction to occur. Such powerful signals may effectively permit “overwriting” of pre-existing neutral place fields with those for salient locations, whereas modest salience signals only facilitate *de novo* place field formation from non-place cells.

### Salience coding in the deep CA1 sublayer

The finding that the time course and magnitude of increases in place cell density vary depending on the nature and extent of salient features advances the idea that the hippocampal deep CA1 sublayer is specialized for salience mapping and proposes a map-based hippocampal salience code that is different from the known firing rate-based coding scheme (Leutgeb 2005b; Komorowski et al., 2009). Unlike previous studies, we observed that the over-representation of salient locations arose in a task that involved learning of salience-place associations through repetitive experience rather than explicit goal-directed spatial navigation (Dupret et al., 2010; Danielson et al., 2016). This observation favors the notion that hippocampal encoding of salience occurs rather automatically without effortful goal-driven learning. Rapid mapping of motivational salience is consistent with its strong behavioral relevance as the source of positive reinforcement, whereas gradual mapping of environmental salience presumably reflects experience-dependent learning of the environment that could contribute to landmark-based navigation (Sato et al., 2017). The extra quantity and stability as well as the idiosyncratic salience-chasing property of place cells for salient locations imply that such “complex” place cells may constitute subpopulations distinct from “simple” place cells for neutral locations in the CA1 deep sublayer (Gauthier and Tank, 2017; Geiller et al.,2017).

The present findings raise a crucial question as to how the presence of salience is signaled to the CA1 deep sublayer. The emergence of reward and landmark overrepresentations with different time courses suggest that signals for each salience arise from different sources, the same source with different patterns and intensities, or perhaps a combination of these. Recent studies demonstrate that reward-responsive ventral tegmental area (VTA) neurons are reactivated during rest periods (Gomperts et al., 2015; Valdés et al., 2015) and optogenetic activation of dopaminergic VTA inputs to CA1 sustains newly acquired hippocampal spatial representations and memory (McNamara et al., 2014), pointing to VTA dopaminergic input as a potential signal for motivational salience. The hippocampus integrates spatial information from the medial entorhinal cortex (MEC) with non-spatial information from LEC to represent objects and events within a spatial context (Knierim et al., 2006; Eichenbaum and Cohen, 2014). LEC neurons start to fire in the vicinity of objects and reward when these features are introduced to the environment (Deshmuk and Knierim, 2011; Tsao et al., 2013). Notably, long-range inhibitory projections from LEC to the dorsal CA1 area respond to diverse salient stimuli, including aversive, motivational and other sensory cues (Basu et al., 2016), making them a candidate for the putative salience signals. Although salience signals may further arise from brain areas other than VTA and LEC, these signals may act to stabilize representations of salient locations in the CA1 deep sublayer, potentially through enhanced reactivation of relevant experience (Singer and Frank, 2009). The deep sublayer-specific CA2 input and mutual suppression between the two sublayers (Kohara et al., 2014; Lee et al., 2014) may help to associate the deep sublayer map more with those salience signals.

## Acknowledgments

We thank Charles Yokoyama, Thomas McHugh, Shigeyoshi Fujisawa and Liset Menendez de la Prida for their comments on the manuscript. This work was supported by Precursory Research for Embryonic Science and Technology (PRESTO) JPMJPR12A1 from the Japan Science and Technology Agency (JST) and KAKENHI Grants 21800091, 24700403, 25116528, 26115530 and 17H05985 from the Ministry of Education, Culture, Sports, Science and Technology (MEXT)/Japan Society for the Promotion of Science (JSPS) to M.S., RIKEN, NIH grant R01DA17310, Grant-in-Aid for Scientific Research on Innovative Area "Foundation of Synapse and Neurocircuit Pathology" from MEXT, Human Frontier Science Program, and High-end Foreign Experts Recruitment Program of Guangdong Province to Y.H., KAKENHI Grants 26870577 to T.T., 25830023 and 15H01571 to K.M., 15H04265 to T.F and 26115504, 25111703 and 21115504 to J.N, and Regional Innovation Cluster Program (City Area Type, Central Saitama Area) from MEXT to J.N. and M.O. D.G-D. is a recipient of the Ph.D. fellowship BES-2013-064171 and a grant from the short-term visit program EEBB-I-15-09552.

## Author Contributions

M.S. and Y.H. designed the study. M.S. and M.K generated the Thy1-G-CaMP7 transgenic mice. M.S. and T.I. built the virtual reality set-up. M.S., K.M., M.K., D.G.-D. and K.K. performed the imaging experiments. M.S., K.M., T.I. and T.T. analyzed the data. M.O. and J.N. made the G-CaMP7-T2A-DsRed2 transgene. T.T., M.S., H.Y. and T.F. developed the image analysis software. M.S. and Y.H. wrote the paper.

## Declaration of interests

Y.H. was supported in part by Takeda Pharmaceutical Co. Ltd. and Fujitsu Laboratories.

## Figure legends

**Figure S1.**
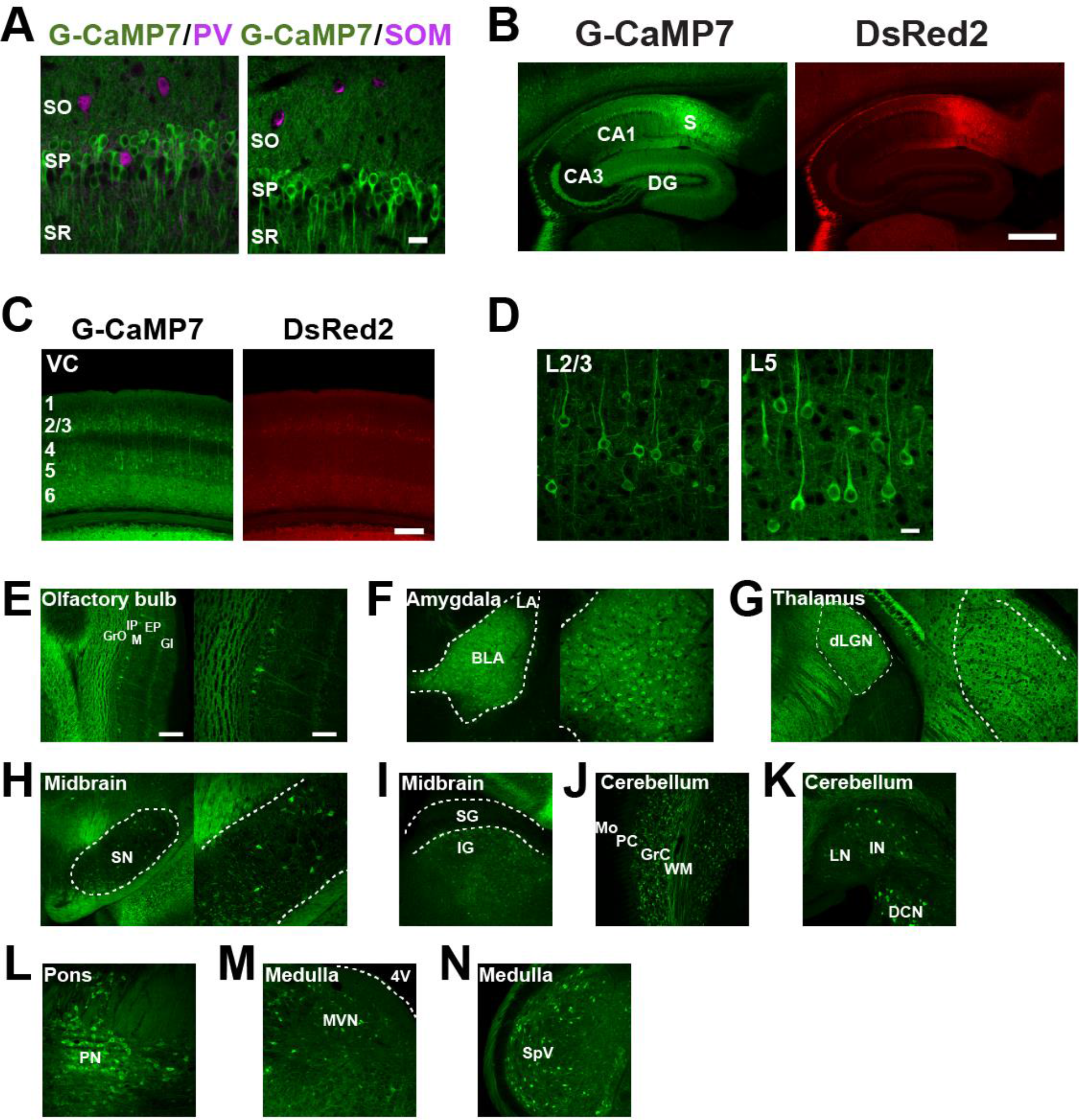
Transgene expression in Thy1-G-CaMP7 transgenic mice. (A) G-CaMP7 expression (green) overlaid with parvalbumin (PV, left) or somatostatin (SOM, right) immunofluorescence (magenta) in the dorsal hippocampal CA1 area at 4 months of age. SO, stratum oriens; SP, stratum pyramidale; SR, stratum radiatum; Scale bar = 20 μm. (B) Expression of G-CaMP7 (left) and DsRed2 (right) in a parasagittal section of the hippocampus at 1 month of age. DG, dentate gyrus; CA1, CA1 area of the hippocampus; CA3, CA3 area of the hippocampus; S, subiculum. Scale bar = 500 μm. (C) Low magnification images of G-CaMP7 (left) and DsRed2 (right) expression in the visual cortex at 3 months of age. Scale bar = 200 μm. (D) High magnification images of G-CaMP7 expression in layer 2/3 (L2/3, left) and layer 5 (L5, right) in the visual cortex at 3 months of age. Scale bar = 20 μm. (E-N) G-CaMP7 expression at 1 month of age in the olfactory bulb (E), amygdala (F), thalamus (G), midbrain (H, I), cerebellum (J, K), pons (L) and medulla (M, N). EP, external plexiform layer; GI, glomerular layer; GrO, granule cell layer of the olfactory bulb; IP, internal plexiform layer; M, mitral cell layer; BLA, basolateral amygdala; LA, lateral amygdala; dLGN; dorsal lateral geniculate nucleus; SN, substantia nigra; IG, intermediate gray layer of the superior colliculus; SG; superficial gray layer of the superior colliculus; GrC, granule cell layer of the cerebellum; Mo, molecular layer; PC, Purkinje cell layer; WM, white matter; DCN, dorsal cochlear nucleus; IN, interposed cerebellar nucleus; LN, lateral cerebellar nucleus; PN, pontine nucleus; MVN, medial vestibular nucleus; 4V, fourth ventricle; SpV, spinal trigeminal nucleus. Scale bar = 200 μm (left panels of E, F, G and H as well as I, L, M and N) or 100 μm (right panels of E, F, G and H as well as J and K).

**Figure S2.**
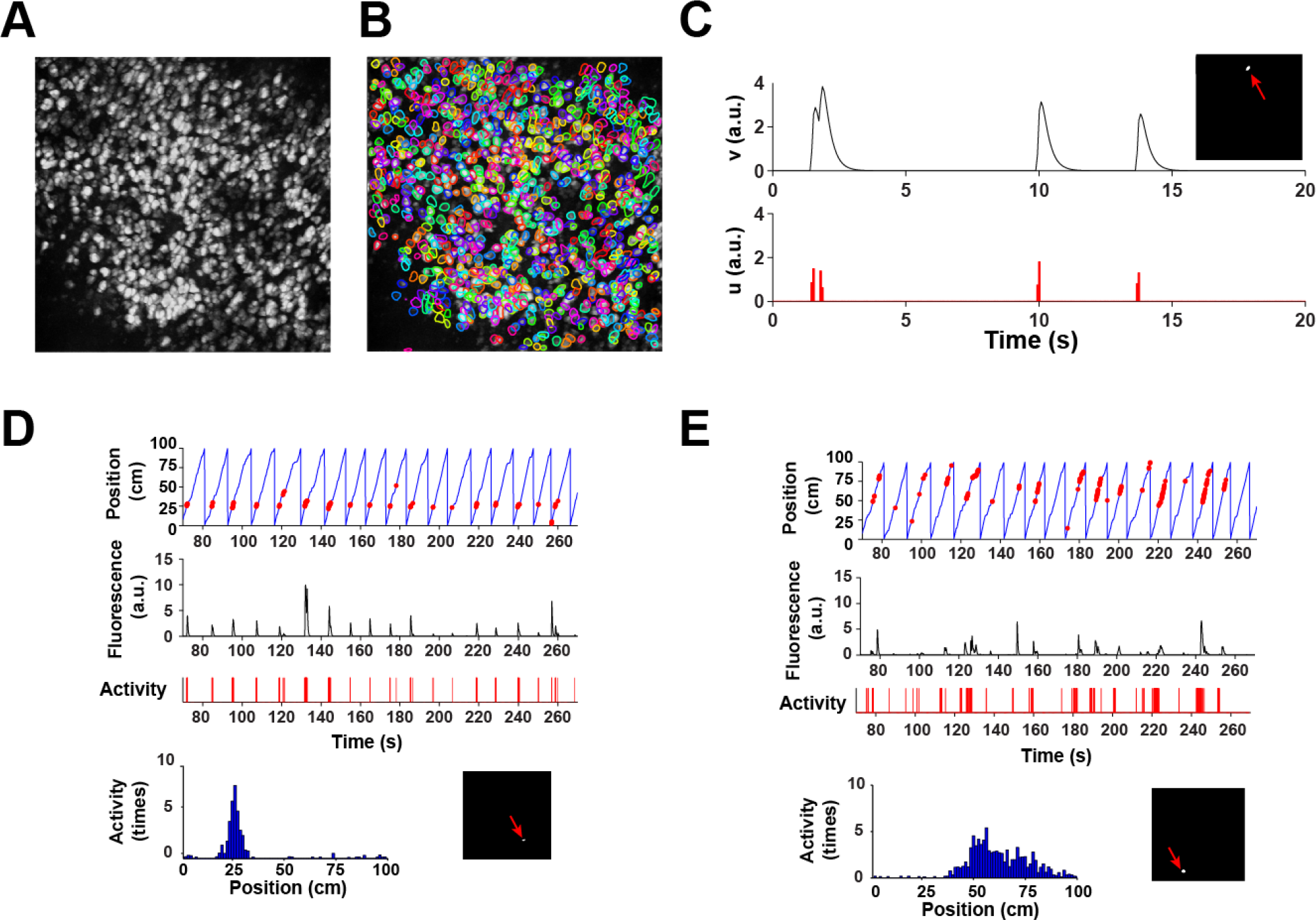
Spatiotemporal deconvolution of calcium imaging data using a non-negative matrix factorization algorithm. (A) A background-subtracted maximum intensity projection image that represents active cells in this example session. (B) Cell identification using the algorithm. In this session, 942 cells were identified. The contours of individual cells are shown by lines of different colors. (C) Time traces of fluorescence intensity (v, top) and inferred spike trains (u, bottom) of an example cell. The anatomical position of the cell is indicated in the inset by a red arrow. (D) Example of vPCs. Shown from top to bottom are a time series of the mouse’s virtual position with the timing of cellular activity indicated by red dots (top), a time series of fluorescence intensity and inferred cellular activity (middle), a histogram of cellular activity plotted against the position in the virtual linear track (bottom left) and the anatomical position of the cell (bottom right). (E) Another example of vPCs. This cell was imaged in a different part of the same field of view as D and activated at a different location in the virtual linear track.

**Figure S3.**
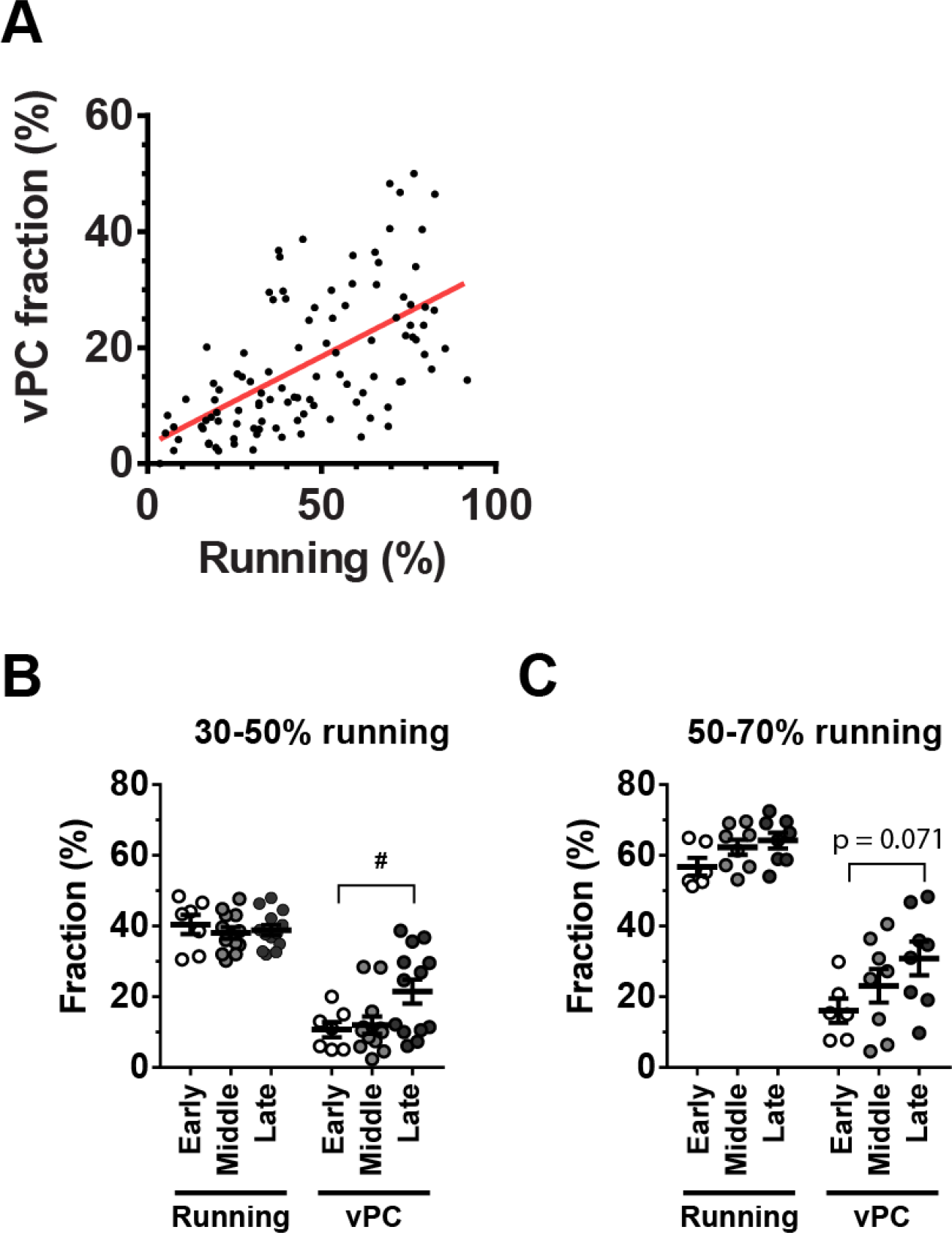
Experience enhances more effective formation of vPCs. (A) The fractions of vPCs were plotted against the corresponding fractions of time spent running for all sessions (n = 105 sessions). The red line represents linear regression (r = 0.59). (B, C) Bar graphs indicating mean fractions of time spent running and those of vPCs in the early, middle and late phases of training. The sessions in which mice ran 30-50% (B) and 50-70% (C) of the time were analyzed. ^#^P = 0.047, F_(2,29)_ = 4.07, n = 7, 12 and 13 sessions, one-way ANOVA. The comparison between Early vPC and Late vPC in 50-70% running exhibited a near-significant trend (P = 0.071, F_(2,19)_ = 2.48, n = 6, 8 and 8 sessions).

**Figure S4.**
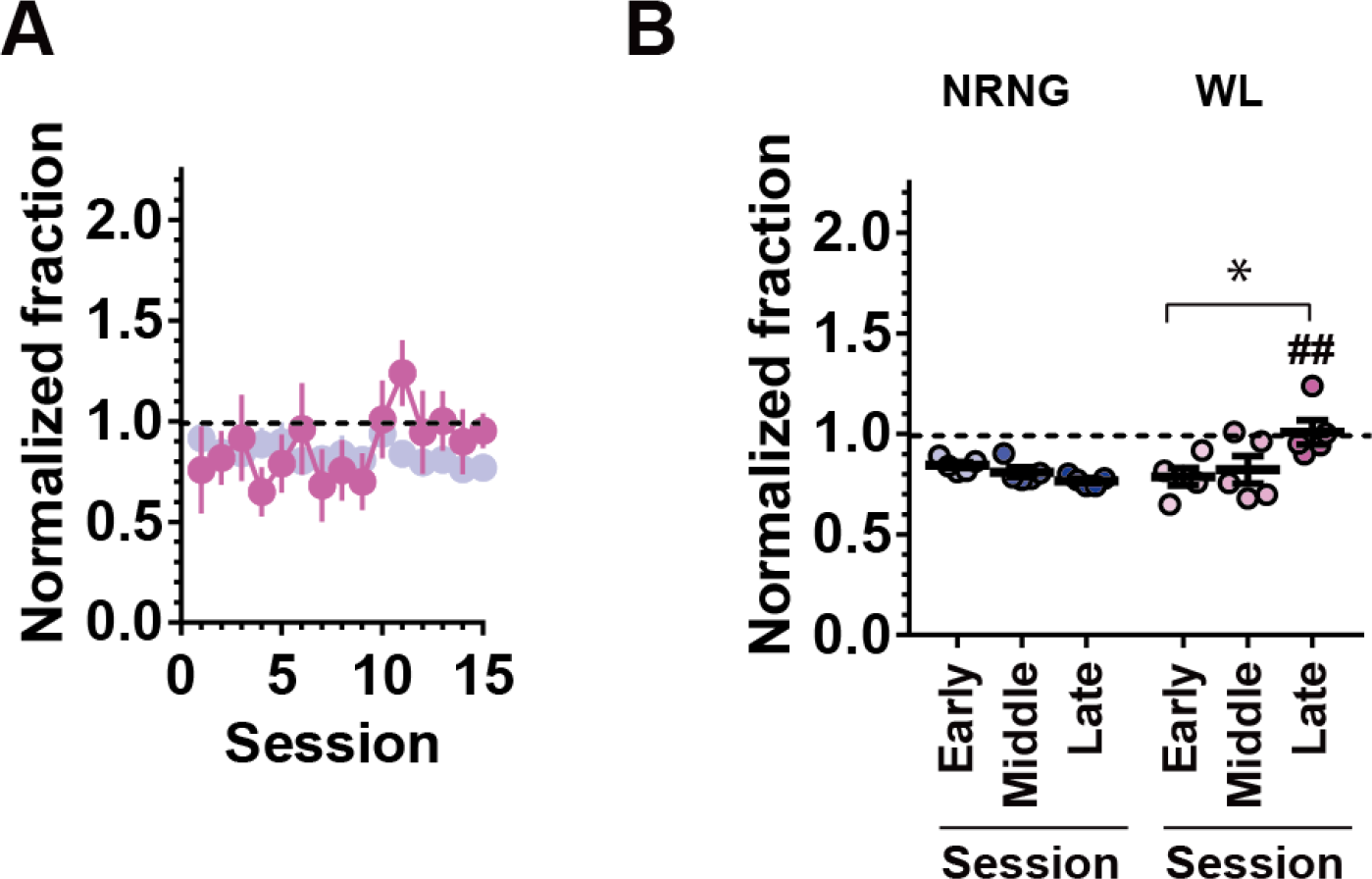
A delayed increase in vPCs that encode a location of a boundary of different wall patterns. (A) The fractions of wall cells (magenta) in each session are expressed as normalized fractions relative to the uniform distribution of vPCs. The data on NRNG in Figure 2H are presented again in this plot for comparison (blue). (B) Average normalized fractions of wall cells (WL) for the early, middle and late phases of the training. The data on NRNG in Figure 2I are presented again in this graph for comparison (blue). *P = 0.037, F_(2,12)_ = 4.79, n = 5, 5 and 5 sessions; one-way ANOVA, ^##^P = 0.0079, U_(5,5)_ = 0, vs. Late NRNG, n = 5 and 5 sessions, two-tailed Mann-Whitney test.

**Figure S5.**
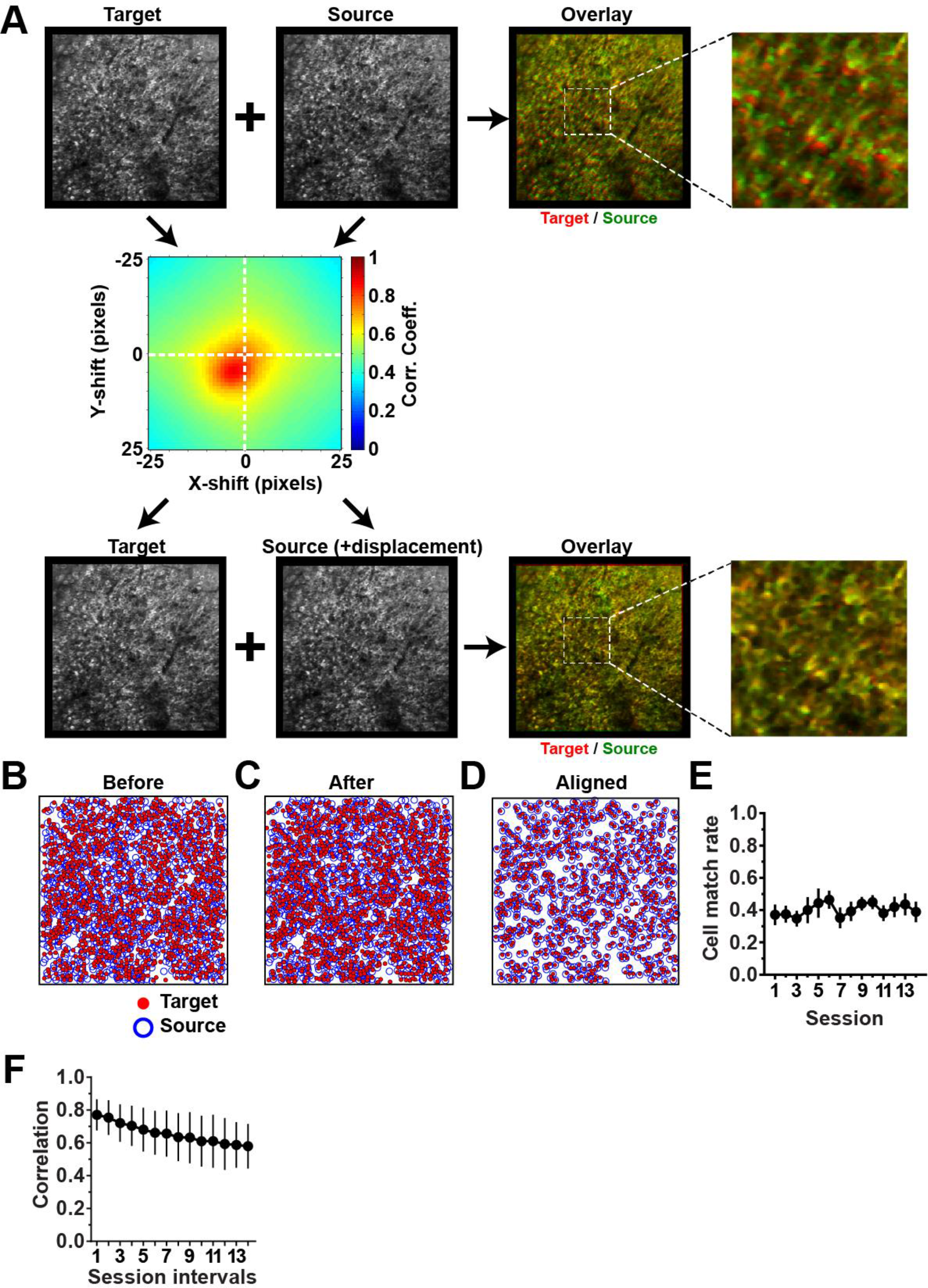
Alignment of cells across sessions. (A) Image registration with or without the displacement estimated by two-dimensional correlation coefficients. A DsRed2 reference image of the target session (top left) was overlaid with that of the source session (top center) without image displacement (top right). This placement resulted in global misalignment of cell positions between the target (red) and source (green) images. To correct this misalignment, two-dimensional correlation coefficients between the two images were calculated within a range of ± 25 × ± 25 pixel displacements (middle). The DsRed2 image of the same target session (bottom left) was then overlaid with that of the source session (bottom center) shifted by the amount of displacement that provided the maximum correlation coefficient (bottom right). This procedure improved global image alignment, as shown by an increase in well-aligned pixels, represented in yellow in the overlaid image. (B) Cells overlaid without correcting image displacement. Cells in the target and source sessions are presented as red dots and blue circles, respectively. (C) Cell aligned after the correction of image displacement. Many cells in both images are now properly aligned, and their local anatomical arrangements are mostly preserved. (D) Cell pairs that were considered to be the same cells are shown (see Methods for detailed criteria). (E) The average fractions of cells aligned between two consecutive sessions. Values are expressed relative to the number of total cells identified in the target sessions. The X-axis indicates the earlier of the two sessions that were compared. (F) The average twodimensional correlation coefficients plotted against the session intervals between the two images compared. Data are presented as the mean ± SD (n = 7 - 98 session pairs).

**Figure S6.**
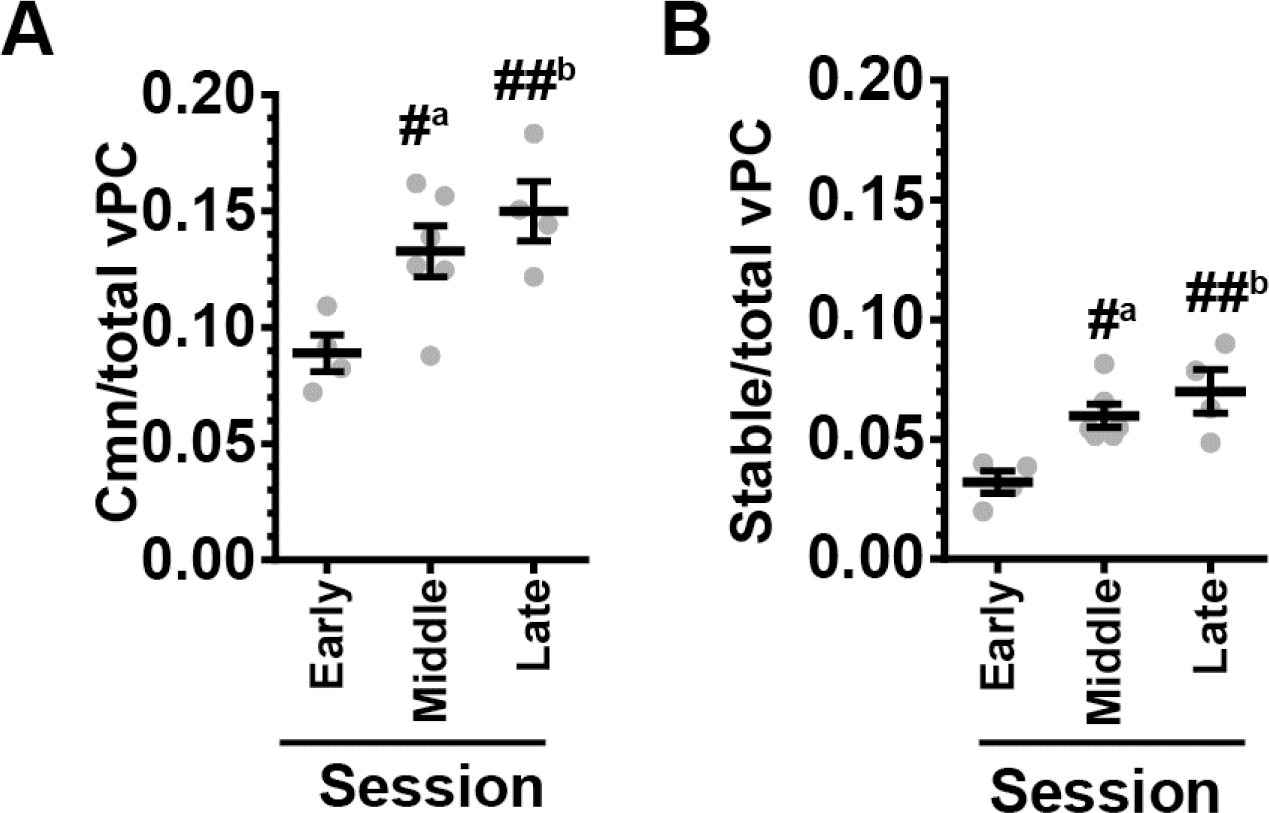
The experience-dependent increase in vPC stability is not due to the increased number of vPCs. (A) Average fractions of common vPCs relative to the number of total vPCs for the early, middle and late phases of training. #^a^, P = 0.029 vs. Early, ##^b^, P = 0.0075 vs. Early, F_(2,11)_ = 7.08; one-way ANOVA, n= 4, 6 and 4 session pairs. (B) Average fractions of stable vPCs relative to the number of total vPCs. #^a^, P = 0.015 vs. Early, ##^b^, P = 0.0037 vs. Early, F_(2,11)_ = 8.90; one-way ANOVA, n= 4, 6 and 4 session pairs.

**Figure S7.**
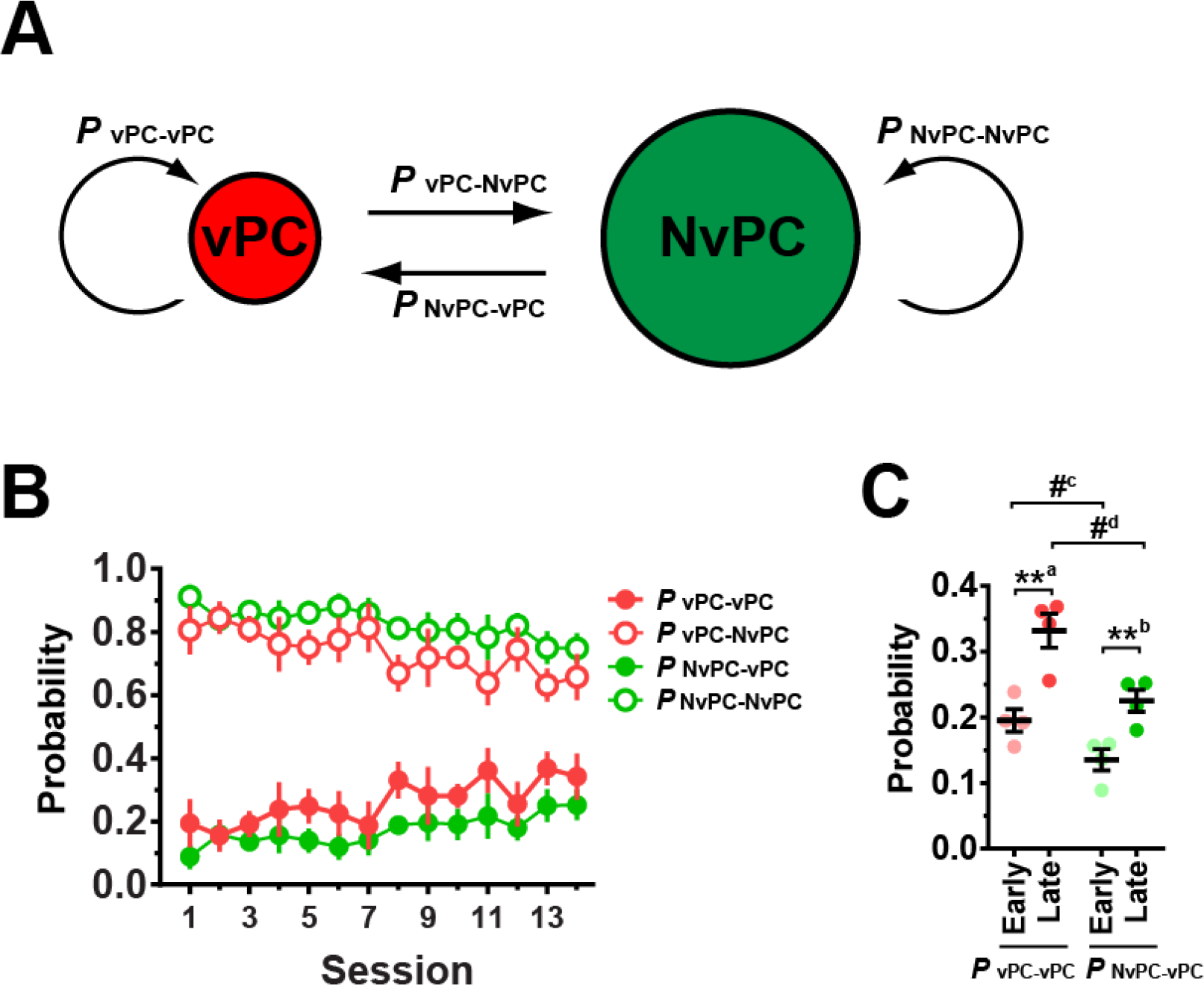
Stability and transition of vPCs and NvPCs across sessions. (A) Two functional cell categories, virtual place cells (vPCs) and non-virtual place cells (NvPCs), and the stability and transition of each category between sessions of interest and their immediately successive sessions are shown in this model. *P* _vPC-vPC_ represents the probability that a vPC in one session remains a vPC in the next session, and *P* _vPC-NvPC_ represents the probability that a vPC in one session becomes an NvPC in the next session. Similar conventions apply to *P* _NvPC-vPC_ and *P* _NvPC-NvPC_. (B) Stability and transition of vPC and NvPC. The X-axis indicates the earlier of the two sessions that were compared. (C) The average probabilities of vPC stability (*P* _vPC-vPC_) and vPC formation (*P* _NvPC-vPC_) for the early and late phases of training. **a P = 0.0046, t_(6)_ = 4.40; **^b^, P = 0.0086, t_(6)_ = 3.84; #^c^, P = 0.044, t_(6)_ = 2.55; #^d^, P = 0.014, t_(6)_ = 3.45; unpaired two-tailed t-test, n = 4 session pairs each.

**Figure S8.**
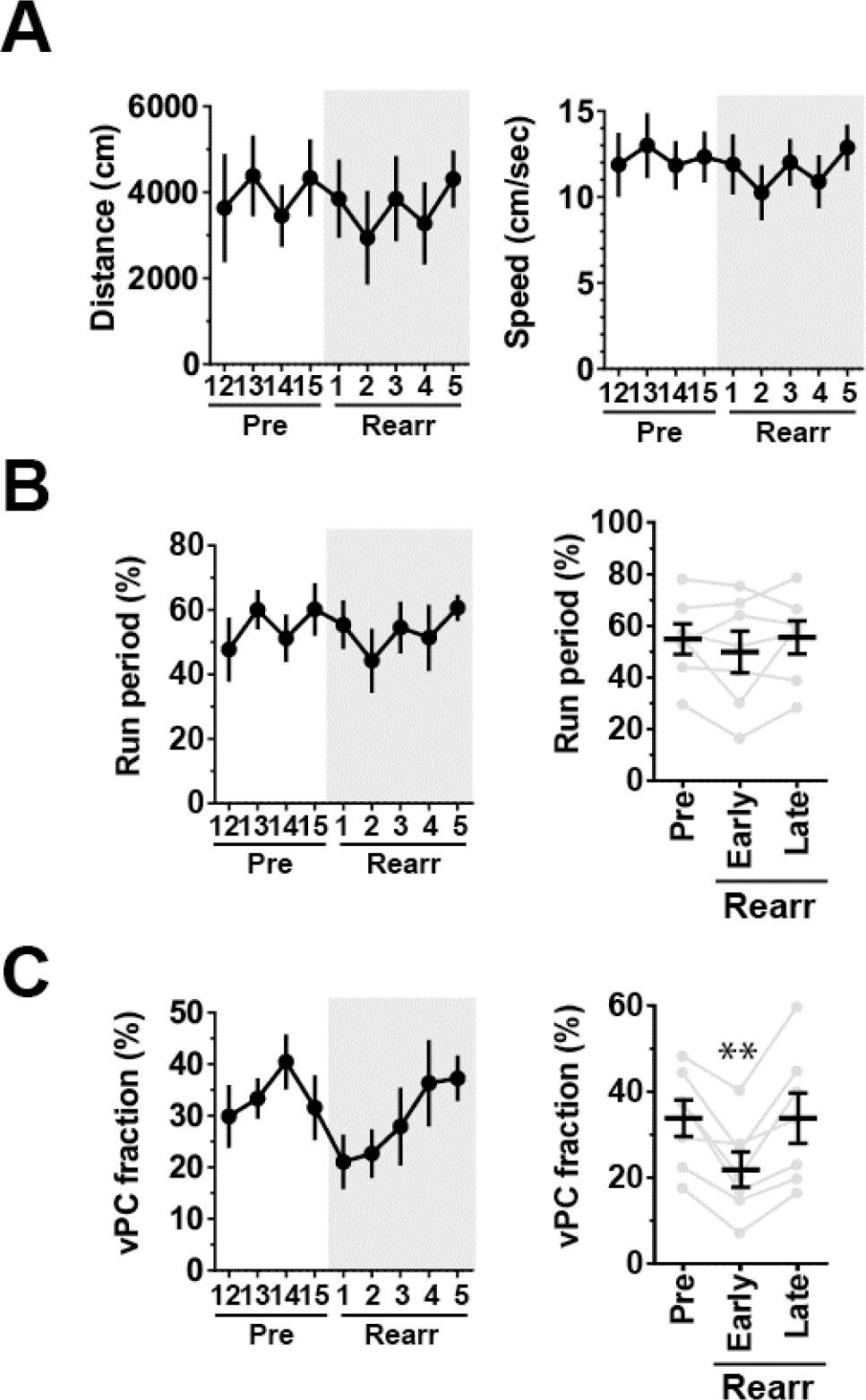
Behavior and vPC fractions during the reward rearrangement task. (A) Total distance traveled (Distance, left) and running speed (Speed, right). (B) The fraction of time spent running (Run period). (C) The fraction of vPCs. The periods during which the reward delivery was relocated are shaded in gray. In B and C, plots indicating average values for pre, early and late phases of re-training are shown on the right. **P = 0.0018 vs. Pre, F_(2,12)_=12.6, one-way ANOVA, n = 7 mice from 2 groups.

**Figure S9.**
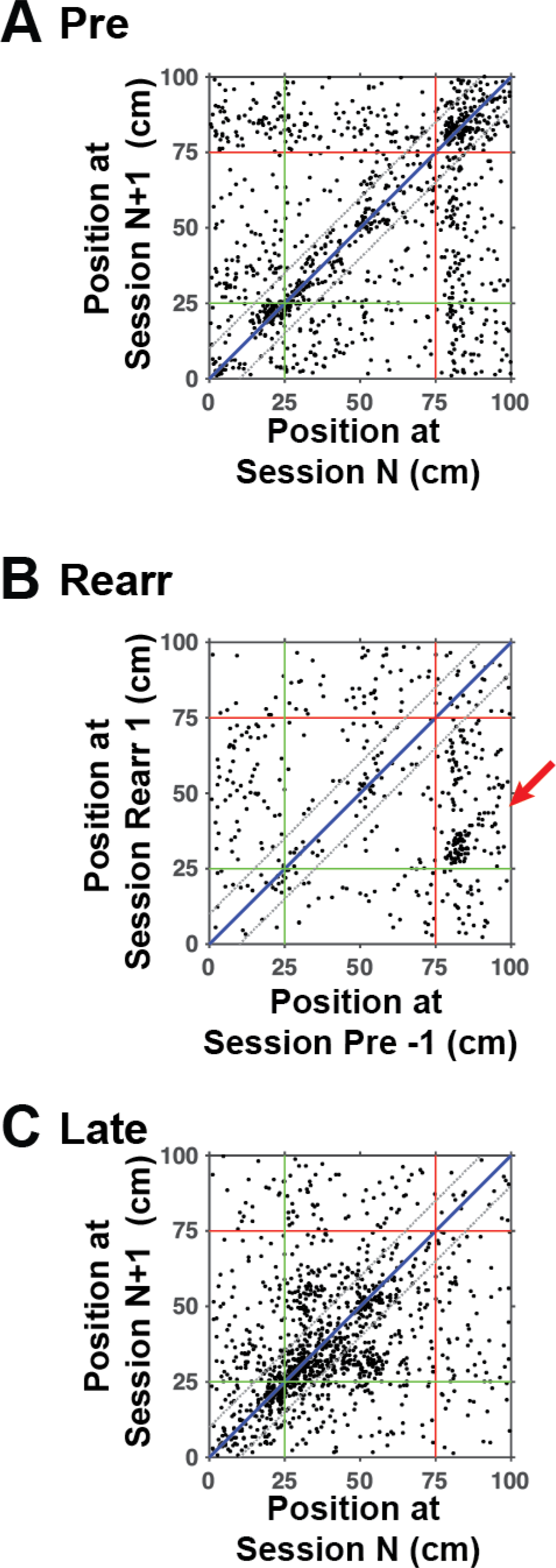
A subpopulation of cells that persistently encode the reward locations. The positions of virtual place fields of common vPCs in two consecutive sessions during the pre-training (A, Pre; n = 1106 cells), before and after the reward rearrangement (B, Rearr; n = 462 cells), and during the late phase of the reward rearrangement task (C, Late; n = 1281 cells) are displayed as two-dimensional plots. The X-axis and Y-axis indicate the positions of the earlier and later of the two sessions, respectively. Each dot represents the position of a cell’s virtual place field, and cells from all mice and relevant sessions are shown. The red arrow in B indicates a cluster of cells that were active at the reward locations regardless of their absolute positions. The green and red lines delineate the positions of the landmark and reward delivery before the reward rearrangement, respectively. The blue diagonal line and the two flanking black dotted lines represent the range that defines stable vPCs.

## Methods

### Ethics statement

All experiments were conducted in accordance with institutional guidelines and protocols approved by the RIKEN Animal Experiments Committee.

### Generation of Thy1-G-CaMP7 transgenic mice

The cDNA encoding G-CaMP7 (Ohkura et al., 2012) ligated to the coding sequence of DsRed2 via a *Thosea asigna* virus-derived 2A peptide sequence (Sato et al., 2015) was subcloned into the Xho I site of the modified mouse Thy-1.2 promoter vector (Feng et al., 2000). The 8.7-kb DNA fragment was prepared by digestion with Not I and Pvu I restriction enzymes and subsequent gel purification and injected into the pronuclei of 466 fertilized eggs of C57BL/6J mice. From 32 offspring, 9 mice were identified as transgene positive, and 6 exhibited transgene expression in the brain. One founder mouse that expressed the transgene at a high level in the hippocampus was used for this study. Mice were genotyped by PCR using the primers 5’-CTGCTGCCCGACAACCA-3’ and 5’-GTCGTCCTTGAAGAAGATGG-3’, which provided a 465-bp product of the G-CaMP7 coding sequence from tail DNA samples of transgene-positive mice.

### Analysis of transgene expression

Thy1-G-CaMP7 transgenic mice were anesthetized deeply with Avertin and perfused transcardially with phosphate-buffered saline (PBS), followed by 4% paraformaldehyde (PFA) in PBS. Brains were removed and further fixed in 4% PFA at 4°C overnight. Parasagittal sections were cut on a vibratome to a thickness of 100 μm. Low-magnification fluorescent images of G-CaMP7 and DsRed2 (Figure 1D) were acquired with a Keyence BZ-9000 epi-fluorescence microscope equipped with a 4x objective. For immunofluorescence labeling, coronal sections were cut on a vibratome to a thickness of 50 μm and incubated with rabbit anti-calbindin D-28K (1:500, AB1778, Millipore, Billerica, MA), rabbit anti-GAD65/67 (1:500, AB1511, Millipore), mouse anti-parvalbumin (1:1000, clone PARV-19, P3088, Sigma, St. Louis, MO) or mouse anti-somatostatin (1:200, clone SOM-018, GTX71935, Gene Tex, Irvine, CA) antibody diluted in PBS containing 2% normal goat serum, 1% BSA, and 0.1% Triton X-100 at 4°C overnight, followed by Alexa 647-labeled goat anti-rabbit or anti-mouse IgG antibody (1:700, A-21245 or A-21236, Thermo Fisher Scientific, Waltham, MA) diluted in the same buffer at room temperature for 1 h. Fluorescence images were obtained using an Olympus FV1000 or FV1200 laser-scanning confocal microscope (Olympus, Tokyo, Japan) equipped with a 20x dry or a 60x water immersion objective lens. The reproducibility of labeling patterns was confirmed in two independent experiments.

### Surgery

Adult male Thy1-G-CaMP7 transgenic mice, at least 12 weeks old and weighing 28-30 g at the beginning of surgery, were used for the experiments. The mice were anesthetized with isoflurane in ambient air (3% induction, 1.5% maintenance) and placed in a custom-made stereotaxic frame. To reduce secretions and brain edema, atropine (0.3 mg/kg, s.c.) and dexamethazone (2 mg/kg, s.c.) were administered prior to anesthesia. A circular piece of scalp was removed, and the underlying bone was cleaned and dried. Three small screws were then placed in the skull (two at the suture of the interparietal and occipital bones and one on the right frontal bone) to provide anchors for the head plate. A thin layer of cyanoacrylate was applied to provide a substrate to which the dental acrylic could adhere.

A stainless steel head plate (25 mm length, 4 mm width, 1 mm thickness) with a wide circular opening (7 mm inner diameter and 10 mm outer diameter, the center is 2.5 mm off relative to the middle of the long side of the plate) was affixed to the skull using dental cement. The center of the opening was targeted at 2 mm posterior to the bregma and 2 mm lateral to the midline in the left hemisphere. The cement was mixed with black ink to block light entry from the LCD monitor into the microscope and placed onto the skull such that it covered the entire skull, including the anchor screws, except for the area of skull inside the opening of the head plate.

Optical window preparation was performed as described previously with modifications (Sato et al., 2016). A few days after the head plate surgery, a 2.5-mm-diameter circular craniotomy was created on the skull overlying the dorsal hippocampus. The dura was removed with forceps, and the overlying cortex was aspirated in a small amount at a time using a blunted 25-gauge needle connected to a vacuum pump. This step was continued with occasional irrigation with cortex buffer (123 mM NaCl, 5 mM KCl, 10 mM glucose, 2 mM CaCl_2_, 2 mM MgCl_2_, 10 mM HEPES, pH 7.4) until the white matter, including the corpus callosum, was exposed. Then, the top-most layers of the white matter were gently peeled aside by holding with the vacuum-connected blunted needle such that its minimal thickness remained covering the dorsal surface of the hippocampus. To minimize bleeding, aspiration was initiated from a cortical area devoid of large vessels, and bleeding was treated immediately with a piece of gelatin sponge (Spongel, Astellas Pharma, Tokyo, Japan) wetted with cortex buffer. An imaging window was then inserted to mechanically support the cranial hole, its surrounding tissue and the hippocampal surface. The imaging window consisted of a stainless steel ring (2.5 mm outer diameter, 2.2 mm inner diameter and 1.0 mm height) with a round coverslip (2.5 mm diameter, 0.17 mm thickness, Matsunami Glass Ind., Osaka, Japan) attached to the bottom using a UV-curable adhesive (NOA81, Norland Products, Cranbury, NJ). To reduce brain movement during imaging, a small disk of medical grade clear silicone sheeting (0.13 mm thickness, 20-10685, Invotec Internartional, Jacksonville, FL) was attached to the surface of the coverslip facing the hippocampal tissue (Mower et al., 2011). When the window was positioned, the bottom coverslip was approximately parallel relative to the head plate, and the hippocampal surface was clearly visible through the bottom coverslip without any trace of bleeding. The upper rim was then cemented to the skull with dental acrylic.

After surgery, a metal cover (0.3 mm thickness) was screwed onto the upper surface of the head plate to protect the imaging window from dust. The mice were placed in a warmed chamber until they fully recovered from anesthesia and were then returned to their home cages. They were housed for at least 4 weeks of postoperative recovery before the start of handling.

### Virtual reality (VR) set-up

A VR system with an air-supported spherical treadmill for head-fixed mice was constructed as described previously (Sato et al., 2017). A 20-cm-diameter Styrofoam ball placed inside the bowl provided a freely rotating surface on which the mouse stood. The mouse was positioned near the top of the ball with its head fixed via the steel head plate that was screwed into a rigid cross bar and posts. A single wide-screen 23” LCD display (Dell U2312, Round Rock, TX) placed 30 cm in front of the mice presented VR scenes rendered by OmegaSpace 3.1 (Solidray Co. Ltd., Yokohama, Japan) running on a Windows 7 computer in 81° horizontal and 51° vertical fields of view. The LCD monitor was large enough to cover the major part of the mouse’s binocular and monocular visual fields (Sato and Stryker, 2008). The use of a single LCD monitor for VR presentation effectively elicits visual cue-based virtual navigation behavior in head-fixed mice (Youngstrom and Strowbridge, 2012; Sato et al., 2017).

The movement of the ball was measured with a USB optical computer mouse (G400, Logitech, Newark, CA) via custom driver and LabVIEW software (National Instruments, Austin, TX). The optical mouse was positioned in front of the mouse and at the intersection of the mouse’s sagittal plane and the equator of the ball. The signals along the horizontal axis (aligned parallel to the mouse’s sagittal plane) generated by the running of the head-fixed mouse was used to compute rotational velocity in the forward-backward direction. This velocity signal was converted into analog control voltages (0-5 V) via a D/A converter and fed to a USB joystick controller (BU0836X, Leo Bodner, Northamptonshire, UK) connected to the OmegaSpace computer to move the mouse’s position in VR.

Water rewards (5 μL/reward) were delivered by a microdispenser unit (O’Hara & Co., Ltd., Tokyo, Japan) attached to a water-feeding tube positioned directly in front of the mouse’s mouth. The unit was triggered upon reward events in VR by 5 V TTL signals generated by an OmegaSpace script via a USB-connected D/A device (USB-6009, National Instruments). The behavioral parameters, such as the mouse’s location in the virtual environment, the trigger signals for water rewards and the rotational velocity signals of the spherical treadmill, were recorded at 20-ms intervals using custom software in LabVIEW. The TTL signals for each frame sent by the microscope computer were recorded with the behavioral data to synchronize the imaging and behavioral data.

### Behavior

At least 5 days before the start of behavioral training, mice implanted with the head plate and the imaging window were acclimated to handling and the Styrofoam ball. During this pre-training session, mice were handled by an experimenter for 5-10 min and then allowed to move freely on the top of the ball, which was rotated manually by the experimenter for another 5-10 min. The procedure was performed once a day and repeated for at least 3 days. The mice were then subjected to a water restriction schedule 2-3 days before start of the behavioral training. Body weight and general appearance were checked daily to ensure that the animals maintained at least ~85% of their pre-surgery body weight and exhibited no signs of abnormal behavior throughout the study. Mice were housed in a group of one to four per cage in 12 h-12 h light-dark cycle (with lights on at 6 pm and off at 6 am on the next day). Experiments were performed during the dark phase of the cycle to enhance the locomotion of the mice.

The virtual endless linear track was created using an editor function of OmegaSpace. The mouse started at the origin of the virtual linear track segment and ran through the track unidirectionally with visual feedback rendered by OmegaSpace. The track segment was 100 cm long, measured as the number of rotations of the ball required to move from one end of the track to the other times the circumference of the ball. The mouse moved only one-dimensionally along the midline of the track with its view angle fixed toward the direction of moving. Different patterns were placed on the walls of each track subsegment as follows: vertical white and black stripes for 0-25 cm; horizontal white and black stripes for 25-50 cm; black dots on a white background for 50-100 cm. The floor was patterned with white grids on a black background. The space above the track was colored black. A green gate was placed as a salient landmark at 25 cm from the origin. Water rewards were delivered when the mouse reached a reward point located 75 cm from the origin. This reward point was located in the middle of a track zone with a certain wall pattern (i.e., black dots on white background) and not denoted with other salient visual cues. Upon reaching the other end of the segment, the mouse’s virtual position was transferred back to the origin, and the same segment of the linear track was presented again. The approaching track segment next to the current one was always rendered on the monitor, so the mouse could see that the virtual linear track was infinitely long.

The mice underwent a total of 15 training sessions in the above task, with 1-2 sessions per day. Each session was 10 min long. When 2 sessions were performed in one day, within-day intervals were at least 4 h, and the mice were returned to their home cages between the sessions. The entire training period from the first to the last sessions was 225 ± 8 h (mean ± SD, n = 7 mice). The mouse was lightly anesthetized with isoflurane to detach the metal window cover screwed onto the head plate and clean the imaging window before being placed into the VR apparatus. The head was then fixed to the crossbar above the ball via the head plate and we waited approximately 20 min until the mouse kept on the ball in darkness recovered fully from the anesthesia. During the behavioral session, the animal was allowed to behave freely in the head-fixed arrangement. G-CaMP7 fluorescence in hippocampal CA1 pyramidal neurons was simultaneously imaged as described below.

For the reward rearrangement task, mice first underwent 15 training sessions in the virtual linear track as described above. The mice were further trained for the following 5 sessions (Rearrangement 1-5) in the same virtual linear track except that the location of reward delivery (75 cm from the origin) was shifted to match the location of the landmark (25 cm from the origin). Data obtained from the last 4 sessions of the initial 15 training sessions before the shift (Sessions 12 through 15, also referred to as Pre −4 through −1) were analyzed as pre-rearrangement baseline sessions. The first rearrangement sessions were performed immediately after the last baseline sessions without releasing the mice from head fixation.

### Imaging

Imaging was performed using a Nikon A1MP (Nikon, Tokyo, Japan) equipped with a 16x, NA 0.8 water immersion objective lens. The microscope was controlled with Nikon NIS-elements software. G-CaMP7 and DsRed2 were excited using a Ti-sapphire laser (MaiTai DeepSee eHP, Spectra-Physics, Santa Clara, CA) at 910 nm. Typical laser power was approximately 40 mW at the objective lens. G-CaMP7 fluorescence was separated using a 560-nm dichroic mirror and collected with an external GaAsP photomultiplier tube (10770PB-40, Hamamatsu Photonics, Hamamatsu, Japan) mounted immediately above the objective lens. The calcium-insensitive DsRed2 fluorescence, which helped to identify G-CaMP7-labeled pyramidal neurons, was simultaneously imaged and recorded using another GaAsP photomultiplier tube. The DsRed2 images were checked by the experimenter for the on-site assessment of the quality of image acquisition but not used for off-line quantitative image analysis, except for image alignment across sessions (Figure S5).

To image G-CaMP7-labeled CA1 pyramidal neurons, the microscope was focused at a depth of approximately 150 μm from the hippocampal surface. To prevent the entry of light from the LCD monitor into the microscope, a small sheet of aluminum foil was wrapped around the objective lens, so the foil completely covered the space between the objective and the skull. Images of 512 × 512 pixels were acquired at a rate of 15 frames per second using a resonant-galvo scanner mounted on the microscope. Each imaging session was 10 min long. The size of the field of view was 532 × 532 μm. In repeated chronic imaging, previously imaged cell populations usually re-appeared at similar depths in new sessions. We took reference images of DsRed2 fluorescence at the beginning of each session to confirm that the reference image of the current session was very similar to that of the previous session by ensuring that blood vessels and neurons arranged in unique patterns appeared in the same parts of the two images.

### Image analysis

Each frame of a G-CaMP7 time-lapse movie was aligned to an average fluorescence image of the movie for motion correction using the TurboReg ImageJ plug-in. The registered movie was then denoised by a spatio-temporal median filter. This preprocessed movie f*(t,x)* was reconstituted to the sum of fluorescence intensity of individual cells using a modified non-negative matrix factorization algorithm, as described in detail elsewhere (Vogelstein et al., 2010; Pnevmatikakis et al., 2016; Takekawa et al., 2017). Briefly, this algorithm assumes that the fluorescence intensity of each cell can be deconvoluted to the spatial filter *a*_*c*_(*x*), which represents the position and shape of the cell, and the time variation *v*_*c*_(*t*) derived from spiking activities *u*_*c*_(*t*):

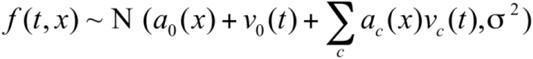

where *a*_*0*_, *v*_*0*_ are baselines, and σ^2^ is intensity of Gaussian noise. As is the case in cell identification using independent component analysis (Mukamel et al., 2009), this algorithm preferentially detects cells that change their fluorescence intensities over time (“active cells”) because cells that barely do so are regarded as being near baseline. Each spike derives the transient elevation of fluorescence intensity with a double-exponential shape:

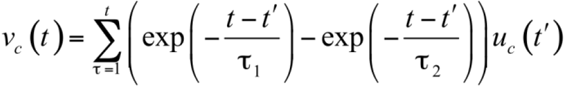

The exponential rise and decay time constants τ_1_ = 0.09 and τ_2_ = 0.261, respectively, were obtained by curve fitting of actual traces of cellular calcium transients in G-CaMP7-expressing CA1 pyramidal neurons in Thy1-G-CaMP7 mice *in vivo*. Spatial filters and spike timings were estimated by two iterative steps. In the first step, we prepared tentative spatial filters and estimated spike trains corresponding to respective filters by a least-squares approach with a non-negative restraint condition. Subsequently, spatial filters were estimated using the least-squares method on the condition that the estimated spike

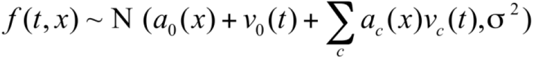

trains were feasible. In addition, we introduced L1 sparse regularization derived from priors that represented the typical cell size and spike frequency. To determine the mutual relationship between a and u, a regularized term was also introduced to the model. This condition guaranteed the uniqueness of the scale of *a*, *u* and *v*. As a consequence, *a*, *u* and *v* are presented in arbitrary units, while the product of *a* and *v* corresponds to the observed data.

In practice, 512 × 512 pixel image data were divided into 4 × 4 of 128 × 128 pixel subareas with 32-pixel overlap regions. Each subarea was analyzed with the above algorithm, and the results were combined to cover the whole image area. After the initial calculation, the morphology of each spatial filter was defined as the region above 0.2 times its peak value, and the position of the filter was defined by its weighted centroid. We then removed the following filters as those that did not represent complete single pyramidal cell morphology: (1) filters whose areas were smaller than 25 pixels, (2) filters whose areas were larger than 400 pixels, (3) filters located on the edge of the image, (4) filters whose heights or widths were greater than 64 pixels because they often contained structures of multiple cells, and (5) smaller filters in filter pairs whose distances were closer than 10 pixels (10.4 μm) and whose temporal correlation coefficients of activities were greater than 0.3 because they were considered to be derived from the same cell.

After those non-cell filters were removed, we recalculated the activity time series for the new filter set. Visual inspection confirmed that nearly all active cells that were represented in a background-subtracted maximum-intensity projection image were identified with this procedure (Figure S2A-B). All images of the entire session, regardless of the mouse’s behavioral state, were used for this image analysis. The average number of cells identified from a movie of a session was 900 ± 246 (mean ± SD, n = 105 sessions).

### Analysis of virtual place fields

Place fields were calculated using cellular activity during movement periods. We defined these periods as the time when the mouse moved at a speed of > 0.5 cm/s continuously for a duration of > 2 s to reject irrelevant movements, such as grooming and jittering on the ball. We divided the entire virtual linear track segment into 80 bins (bin size = 1.25 cm) and created a histogram of neuronal activity versus track position for each cell. The activity events were defined by binarizing the time series of inferred spike activity *u* at a threshold of 0.1, which was empirically determined to remove baseline noise. The counts of the histogram were then divided by the mouse’s occupancy time at each bin, and the resultant place fields were Gaussian-smoothed (Gaussian window size = 6.25 cm) and normalized to the maximum values. To test the significance of virtual place-related activity, we calculated the mutual information content between neuronal activity and the mouse’s virtual location for each cell (Markus et al., 1994; Ziv et al., 2013). We compared this value to a distribution of mutual information content calculated using 1000 randomly permuted data for the same cell. The permutation was conducted by rotating the activity event time series by a random amount relative to the time series of the mouse’s virtual positions. Cells were considered to be virtual place cells (vPCs) if their overall activity rates within the session were no less than 0.1 events/s and their mutual information contents in the real data were greater than the 95th percentile of the values obtained from the randomly permuted data. We defined the position of the virtual place field of each vPC by the position of the peak of the field. A vPC was considered to be a “gate cell”, “reward cell” or “wall cell” if its virtual place field position was 17.5-32.5, 75-95 or 47.5-55 cm from the origin of the track segment. vPCs with virtual place field positions outside the above zones were categorized as “non-reward, non-gate vPCs.” The vPC formation factor was defined by the slope of a least-squares regression line fitted to a plot of the fraction of vPCs against the fraction of time spent running, which contained data points from all animals in the session of interest. The linear regression model included no constant term under the assumption that no vPCs were formed without running in each session. When calculating the fractions of gate cells, reward cells and non-reward non-gate vPCs relative to the number of total vPCs, data from sessions with at least 35 total vPCs (n = 93 sessions) were used to avoid the effects of improperly large or small fractions caused by small cell numbers.

### Bayesian decoding

Bayesian probability-based reconstruction of the subject’s trajectory from imaging data was conducted essentially as described by Zhang et al.(Zhang et al., 1998; Ziv et al., 2013). We computed P(**x**|**n**), the conditional probability for the subject to be at location **x** given the neuronal activity **n** occurred within a time window as follows. P(**x**), the unconditional probability for the subject to be at position **x** was calculated from the dwell time distribution at each spatial bin (bin size = 1.25 cm). P(**n**|**x**), the conditional probability for neuronal activity **n** to occur given the subject is at position **x**, was computed using *f*_*i*_(**x**), the spatial map of neuronal activity of cell *i* at position **x,** under the assumption that occurrence of each Ca^2^ transient is statistically independent as well as that vPCs are independent of each other (Zhang et al., 1998; Ziv et al., 2013). Time-varying activity traces containing Ca^2^ transients were transformed into binarized vectors of the neuronal activity (time step = 67 ms), in which 1 and 0 represent the presence and absence of a Ca^2^ transient, respectively, and these vectors were binned in 200 ms bins for probability calculations. P(**n**), the probability for neuronal activity **n** to occur, was estimated by normalizing P(**x**|**n**) by the condition that the sum of P(**x**|**n**) along **x** is equal to 1 (Zhang et al., 1998). The peak position of the resultant probability distribution of P(**x**|**n**), computed using the above conditional and unconditional probabilities by Bayes’ theorem, was regarded as the reconstructed position of the subject and the entire trajectory was obtained by sliding the time window ahead. Estimation error was calculated as the difference between the real and the reconstructed positions. Data from the sessions with a running time ≧ 240 s were used for the reconstruction (n= 57 sessions). We trained the decoder with the subject’s observed virtual positions and activities of all vPCs during the first 150 s of the running period and estimated the trajectory for the following 90 s running time of the same sessions using the corresponding vPC activities. The sessions with average median errors across all positions less than 10 cm were classified as well-decoded sessions (n= 25 sessions), and the errors of these sessions were then averaged separately for the locations encoded by GT, NRNG, and RW cells for comparison.

### Alignment of cells across sessions and analysis of cell transitions

To find a population of the same cells in images that were acquired in two different sessions, we first estimated the extent of overall image displacement that existed between the two image datasets. We searched for a peak in the two-dimensional correlation coefficient calculated between the two DsRed2 reference images obtained at the beginning of each session within a range of a 25 × 25 pixel (26.0 × 26.0 μm) displacement in the x and y dimensions (Figure S5A). All compared image pairs displayed a peak within this range (average displacement in the x dimension, 5.9 ± 4.5 μm; average displacement in the y dimension, 4.7 ± 4.2 μm; average peak correlation coefficient 0.77 ± 0.09, mean ± SD, n = 98 image pairs). During the calculation of two-dimensional correlation coefficients, the image of one session (the “source” session) was systematically shifted relative to that of the other session (the “target” session). The map of the coordinates of all cell positions in the target session was then overlaid with that of the source session, shifted by the amount of the estimated displacement (Figure S5A). The cell closest to each cell in the target session was searched in the displaced source session map, and the cell that was found was regarded provisionally to be the same cell if they were separated by 5 pixels (5.2 μm) or less. Cells that were unable to find the closest cells within this range were rejected from the subsequent analysis. After finding the provisional counterparts in the displaced source session map, the same procedure was repeated for the cells in the displaced source session map to conversely find their closest partners in the target session. This step helped remove cell pairs that were redundantly assigned (e.g., two different cells in one session falsely assigned to the same single cell in the other session) and the resultant cell pairs that had mutually unique correspondence were considered to be the pairs that represented the same cells (termed hereafter “common cells”). When comparing vPC maps, common vPCs were defined as a subset of common cells that were identified as significant vPCs (see above) in both consecutive sessions. Stable vPCs were defined as a subset of common vPCs with virtual place field positions in the consecutive sessions that were close to each other (i.e., virtual place field distance < 10 cm). The stability and transition probabilities of vPCs and non-vPCs (NvPCs) (Figure S7) were calculated as follows. First, we identified a population of common cells that belonged to the cell category of interest in the reference session N and then examined the category into which each of them was classified in the subsequent session N+1. The probability of transition from category A in session N to category B in session N+1, *P* _A-B_, was calculated by dividing the number of cells that belonged to category A in session N and became a cell of category B in session N+1 by the total number of cells that belonged to category A in session N. The probability of stability was calculated similarly by considering it as a process of transition to the same category. The analysis of formation, recruitment and stabilization of vPCs (Figures 5 and 8) was conducted similarly, except that cells were classified into four categories in the reference session N, and the position of the virtual place field of each cell was tracked in the subsequent session N+1. The results are expressed as the cell density rather than probabilities to show the relative contributions of the different cell categories.

### Statistics

Data are expressed as means ± SEM unless stated otherwise. When only two groups were compared, two-sided Student’s t-tests were used if the variances of the two groups were similar. Otherwise, two-tailed Mann-Whitney tests were used. When more than two groups were compared, analysis of variance (ANOVA) was used if variances of the groups compared were similar. Otherwise, a non-parametric version of ANOVA (Friedman test) was used. In both parametric and non-parametric ANOVA, P-values were adjusted for post hoc multiple comparisons. Exact P-values are shown unless P < 0.0001. Statistical tests were performed using GraphPad Prism Version 6.05 (GraphPad Software, Inc., La Jolla, CA).

